# Insights into Intestinal P-glycoprotein Function using Talinolol: A PBPK Modeling Approach

**DOI:** 10.1101/2023.11.21.568168

**Authors:** Beatrice Stemmer Mallol, Jan Grzegorzewski, Hans-Michael Tautenhahn, Matthias König

## Abstract

Talinolol is a cardioselective beta-blocker that was previously used to treat heart failure and myocardial infarction. Following the development of new, more effective beta-blockers with better study results, talinolol is now only used clinically for the treatment of arterial hypertension. In basic science, talinolol continues to be used as a test substance due to its pharmacokinetics. Its intestinal absorption is determined by uptake by the organic anion transporting polypeptide 2B1 (OATP2B1) and efflux via P-glycoprotein (P-gp). Talinolol can be taken up via OATP1B1 in the liver, where it enters the enterohepatic circulation. Talinolol is excreted unchanged in the urine and feces. Talinolol is widely used as a probe drug for the intestinal efflux transporter P-gp, which plays a critical role in protecting against potentially toxic substances and facilitating the elimination of xenobiotics. In this work, an extensive database of talinolol pharmacokinetics was established and used to develop and validate a physiologically based pharmacokinetic (PBPK) model of talinolol for P-gp phenotyping. The model was used to investigate the influence of several factors on talinolol pharmacokinetics: (i) inhibition of P-gp via drug-drug interaction; (i) genetic polymorphisms of P-gp; (iii) activity of OATP2B1 and OATP1B1; (iv) effect of comorbidity, namely hepatic and renal impairment; and (v) site-specific distribution of P-gp and OATP2B1 in the intestine. The model accurately predicts the concentration-time profile of talinolol after oral or intravenous administration of single and multiple dosing. Furthermore, the model accurately describes the effect of genetic variants of P-gp on the pharmacokinetics of talinolol, the effect of inhibition of P-gp, the effect of renal impairment, as well as site-specific infusion of talinolol in the intestine. The detailed description of the intestinal absorption of talinolol and the predictions of talinolol pharmacokinetics as a function of hepatorenal impairment provide valuable clinical insights for metabolic phenotyping with talinolol. Both the model and the database are freely available for reuse.

## 1 INTRODUCTION

Talinolol is a cardioselective beta-blocker of the β1 adrenergic receptor. As such, it helps to reduce heart rate, cardiac output, and blood pressure, making it useful in the treatment of various cardiovascular conditions, particularly arterial hypertension. It’s most notable for its interaction with the intestinal efflux transporter P-glycoprotein (P-gp), which affects its absorption and pharmacokinetics. This specific interaction has led to the use of talinolol as a probe drug to study the function of P-gp in the intestine. The drug is usually administered orally, but there is also an option for intravenous application.

P-glycoprotein (P-gp), also known as multidrug resistance protein 1 (MDR1), is a major membrane protein belonging to the ATP-binding cassette (ABC) transporter family and is encoded by the ABCB1 gene (Cascorbi, 2006). It is widely expressed in various tissues, particularly in intestinal epithelium, liver, kidney, and the blood-brain barrier (Fromm, 2004; Thiebaut et al., 1987). P-gp plays a vital role in the body’s defense mechanism by acting as an efflux pump to remove potentially toxic compounds, including many drugs, from cells. This protective function can affect the absorption, distribution, metabolism, and excretion of many drugs. For example, P-gp transports talinolol outwards of the intestinal enterocytes into the small intestine (Gramatté et al., 1996).

The pharmacological properties of talinolol include low plasma protein binding (Tubic et al., 2006b), minor enterohepatic recirculation (Terhaag et al., 1989; Wetterich et al., 1996), and no significant first pass metabolism (Trausch et al., 1995) making it an ideal test substance to study P-gp.

In the intestine, talinolol is taken up by the OATP2B1 transporter across the apical membrane into the enterocytes, from where it can be further transported into the blood. In contrast, P-gp acts as an efflux transporter of talinolol, which exports a fraction of the absorbed talinolol from the enterocytes back into the intestinal lumen. The interplay between OATP2B1 and P-gp influences the amount of talinolol absorbed into the circulation, resulting in a reduced oral bioavailability of 55-75 % for talinolol (Trausch et al., 1995; Gramatté et al., 1996; Giessmann et al., 2004; Bernsdorf et al., 2006; Schwarz et al., 2007). Genetic variants and drug-drug interactions can affect the activity of these transporters. For example, rifampicin and St. Johns wort have been identified as substrates of the pregnane X receptor (PXR), which positively affects P-gp expression (Banerjee and Chen, 2013; Dürr et al., 2000; Haslam et al., 2008). An example of a drug-drug interaction is erythromycin, which inhibits P-gp activity (Schwarz et al., 2000).

Talinolol enters the liver via the uptake transporter OATP1B1 (Bernsdorf et al., 2006). Approximately 10 % of the administered talinolol passes through the enterohepatic circulation from the liver via the bile back into the intestinal lumen (Terhaag et al., 1989; Wetterich et al., 1996; Haustein and Fritzsche, 1981). After oral administration, talinolol is eliminated from the body by urinary excretion, which accounts for approximately 60 %, and by fecal excretion, which accounts for approximately 40 % (Trausch et al., 1995; Tubic et al., 2006b), due to incomplete intestinal absorption and reabsorption.

An important factor in intestinal absorption is the spatial distribution of uptake and efflux transporters across different segments of the intestine, from the duodenum, through the jejunum and ileum, to the colon. The orchestrated interplay of these transporters across these regions is a key determinant of absorption dynamics. While mRNA expression and protein levels of OATP2B1 are constant throughout the small intestine, there is a marked increase in both mRNA expression and protein levels of P-gp from the upper to the lower sections of the intestine (Drozdzik et al., 2014). Notably, studies have identified a direct correlation between duodenal P-gp mRNA expression and the bioavailability of orally administered talinolol (Bernsdorf et al., 2006; Bogman et al., 2005; Schwarz et al., 2007). The use of site-specific infusions offers a refined approach to study regional absorption patterns within the intestine. For example, the results of (Gramatté et al., 1996; Bogman et al., 2005) support the hypothesis of a regioselective absorption window for talinolol.

Genetic polymorphisms that alter the enzyme activity of P-gp and the OATP isoforms OATP2B1 and OATP1B1 could have a major effect on talinolol pharmacokinetics, but there is limited information on the effect of polymorphisms of these transporters on talinolol pharmacokinetics.

P-gp is encoded by the ABCB1 gene, which is known to carry several single nucleotide polymorphisms (SNPs). Notable SNPs are observed in exon 12 (1236C>T), exon 21 (2677G>T/A) and especially 3435C>T exon 26 (Hoffmeyer et al., 2000; Marzolini et al., 2004). The 3435C>T and 1236C>T polymorphisms are synonymous, i.e. they do not cause a change in the corresponding amino acid sequence. In contrast, the 2677G>T/A polymorphism results in the replacement of serine at position 893 by either threonine or alanine (Ambudkar et al., 2003). Most studies show that an increased presence of SNPs in exons 12, 21 or 26 correlates with a reduced area under the curve (AUC) for talinolol (He et al., 2012; Schwarz et al., 2007). However, some studies contradict this and suggest that the 3435C>T polymorphism and the combined 2677G>T/A and 3435C>T variants have no discernible effect on the pharmacokinetics of talinolol (Bernsdorf et al., 2006; Bogman et al., 2005; Han et al., 2009; Siegmund et al., 2002; Zhang et al., 2005).

OATP2B1 and OATP1B1, encoded by the genes SLCO2B1 and SLCO1B1 respectively, are members of the organic anion transporter protein family (Tamai et al., 2000; Abe et al., 1999). While OATP2B1 primarily facilitates drug absorption in the small intestine, OATP1B1 is crucial for hepatic basolateral uptake (Nakanishi and Tamai, 2012). In OATP2B1, the SNPs C1457C>T and G935G>A lead to amino acid changes: Ser486Phe and Arg312Gln respectively. There is little literature on how OATP2B1 polymorphisms affect talinolol absorption. However, with alternative substrates such as fexofenadine (Imanaga et al., 2011) and the beta-blocker celiprolol (Ieiri et al., 2012), the genetic variant SLCO2B1*3 (marked by the c.1457C>T mutation) showed reduced transport activity compared with wild-type SLCO2B1*1, whereas others reported the opposite (Akamine et al., 2010).

OATP1B1 has notable SNPs, namely 388A>G and 521T>C, resulting in amino acid changes from Asn to Asp and Val to Ala, respectively. Important alleles are the wild-type *1a (388A-521T), *1b (388G-521T), *5 (388A-521C) and *15 (388G-521C) (Smith et al., 2005; Tirona et al., 2001). Notably, individuals carrying the SLCO1B1*1b allele show increased protein activity and fecal excretion compared to those with the wild-type SLCO1B11a/*1a genotype. As a result, talinolol is more readily absorbed by the liver in carriers of the *1b allele, resulting in a significantly reduced half-life compared to the wild type (Bernsdorf et al., 2006). In contrast, *5 and *15 show reduced activity.

The aim of this study was to develop a physiologically based pharmacokinetic (PBPK) model to investigate the influence of several factors on the pharmacokinetics of talinolol and the use of talinolol as a probe drug for P-gp: (i) genetic variants of P-gp; (ii) enzymatic activity of the transporters OATP2B1 and OATP1B1; (iii) site-specific distribution of P-gp and OATP2B1 proteins in the intestine; and (iv) effects of concomitant diseases such as liver cirrhosis or chronic renal failure. The overall goal is to better understand how these factors affect the pharmacokinetics of talinolol and contribute to the metabolic phenotyping via talinolol.

## 2 MATERIAL AND METHODS

### Database of talinolol pharmacokinetics

Talinolol pharmacokinetic data were systematically curated from the literature for model development, parameterisation and validation. An initial literature search was performed on PubMed using the query ‘talinolol AND pharmacokinetics’. The literature corpus was expanded with additional references found in the primary sources. In addition, we performed a search using PKPDAI for ‘talinolol’ and merged the literature. We then screened and filtered the resulting publications based on the presence of pharmacokinetic parameters for time course data related to talinolol. Our primary focus was on data from healthy volunteers, but we also included data from patients with kidney or liver dysfunction. In addition, studies that provided information on P-gp, OATP1B1 and OATP2B1 genotypes and associated mRNA or protein data were of particular interest.

The selected studies were reviewed to extract information on various aspects such as subject characteristics, group demographics (including age, sex, specific diseases, medications and genotypes), talinolol dosing protocol and talinolol pharmacokinetic profile. These details were manually curated using established data curation protocols designed for pharmacokinetic information. This process involved digitizing data from figures, tables and textual descriptions as described in (Grzegorzewski et al., 2021). An overview of the 33 curated studies is provided in Tab. 1 with an overview of the literature research in Fig. S1. The extensive heterogeneous data set provided the data base for model development and validation. All data are available in the open database PK-DB (https://pk-db.com) (Grzegorzewski et al., 2021).

**Table 1:**
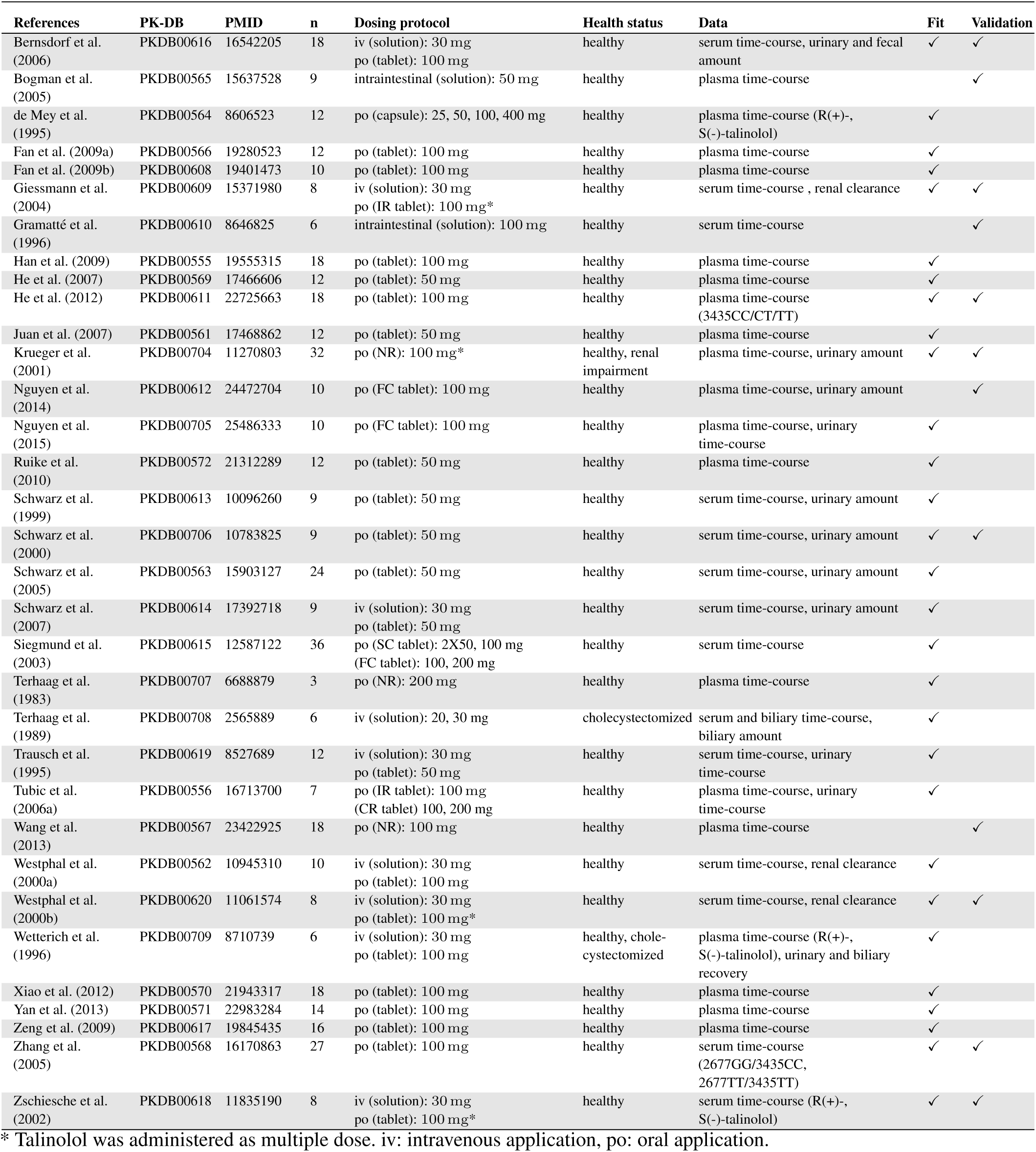
Studies used for the parameterization and validation of the model.

### Model

The PBPK model was encoded in the Systems Biology Markup Language (SBML) (Hucka et al., 2019; Keating et al., 2020). For model development and visualization, sbmlutils (Kö nig, 2021b) and cy3sbml (König et al., 2012; Kö nig and Rodriguez, 2019) were used. The model is based on ordinary differential equations (ODEs), which are numerically solved using sbmlsim (König, 2021a) based on the high-performance SBML simulator libroadrunner (Somogyi et al., 2015; Welsh et al., 2022). The model is made available in SBML under a CC-BY 4.0 license from https://github.com/ `matthiaskoenig/talinolol-model` with model equations available from the repository. Within this work, version 0.9.4 of the model was used (Stemmer Mallol and König, 2023).

The developed PBPK model consists of a whole-body model connecting different organs via the systemic circulation (Fig. 1). The values for organ volumes and tissue perfusion are provided in Tab. 2, parameters for the site-specific intestine model in Tab. 3. The PBPK model follows a hierarchical structure, with the whole-body model connecting the submodels for the intestine, kidneys and liver. The biochemical reactions within the model describe the import and export of talinolol between the plasma and organ compartments. The intestinal import process, mediated by OATP2B1, was modeled through irreversible first-order Michaelis-Menten kinetics, while talinolol export reactions follow irreversible mass-action kinetics. Hepatic impairment was modeled as previously described (Köller et al., 2021a,b). Renal impairment was modeled as a stepwise decrease in renal function by scaling all renal processes with the factor `f renal function`, where 1.0 represents normal function and 0.0 represents no renal function.

**Figure 1:**
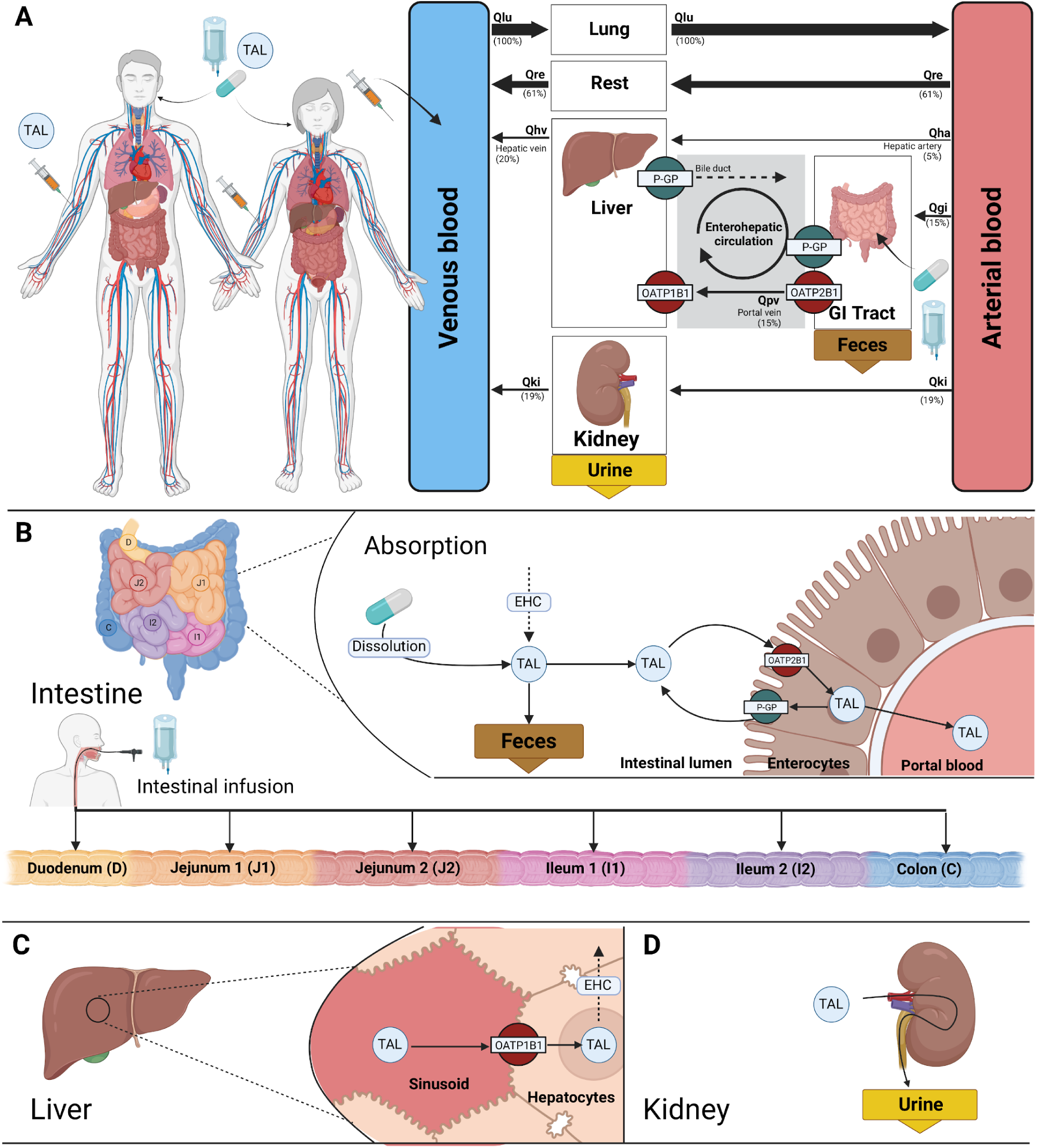
Physiologically based pharmacokinetic (PBPK) model of Talinolol. **(A)** The whole-body model consists of the systemic circulation connecting the organs by blood flow. The gastrointestinal tract, liver, kidneys, and lungs were modeled in detail, with the remaining organs grouped in the rest compartment. Talinolol can be administered either intravenously or orally. **(B)** Intestinal model consisting of dissolution, absorption and fecal elimination. Talinolol can be absorbed from the intestine via OATP2B1 with the efflux transporter P-gp reducing absorption. To model regiospecific intestinal infusion of talinolol, the intestine was divided into the segments duodenum, jejunum 1, jejunum 2, ileum 1, ileum 2 and colon. Within each segment, the model includes the uptake and efflux of talinolol by OATP2B1 and P-gp. Talinolol which is not absorbed is excreted in the feces. **(C)** In the liver model, talinolol is taken up from the blood into hepatocytes via OATP1B1 and subsequently excreted in the bile, shown as the dashed arrow. Via the enterohepatic circulation (EHC), talinolol can be transported from the bile into the intestine. **(D)** Kindey model consisting of urinary excretion of talinolol. Created with Biorender.

**Table 2:**
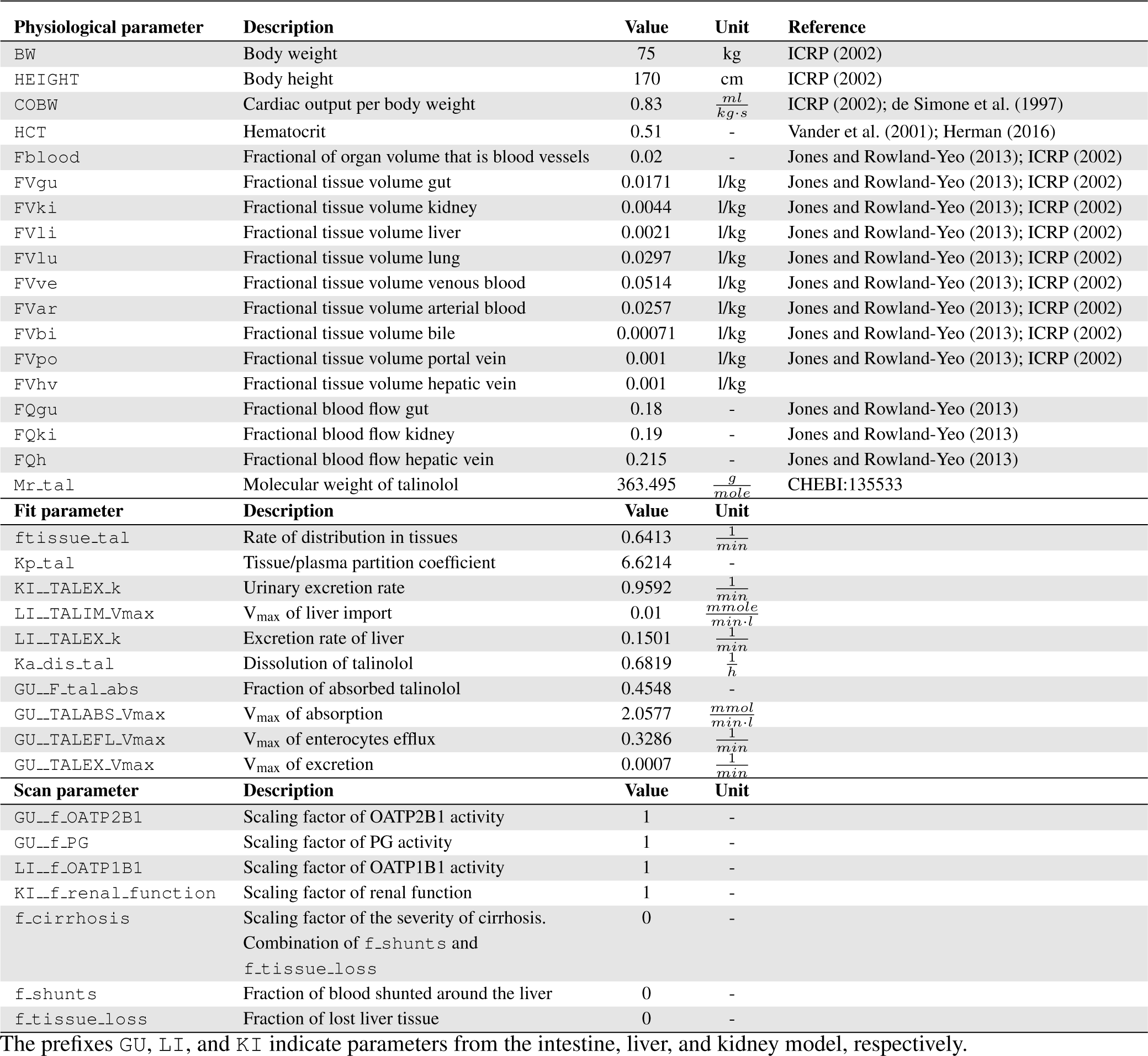
Model parameters.

**Table 3:** Site-specific intestinal parameters.

| Parameter | Duodenum | Jejunum.1 | Jejunum.2 | Ileum.1 | Ileum.2 | Colon | Reference |
| --- | --- | --- | --- | --- | --- | --- | --- |
| Length [cm] | 30 | 105 | 105 | 165 | 165 | 135 |  |
| Diameter [cm] | 3.7 | 2.7 | 2.7 | 2.7 | 2.7 | 6.0 |  |
| Velocity [$\frac{cm}{min}$] | 2.0 | 2.0 | 2.0 | 2.0 | 2.0 | 0.5 | |
| Protein amount P-gp [-] | 0.3 | 0.4 | 0.5 | 0.7 | 1.1 | 0.25 | Englund et al. (2006); Drozdzik et al. (2014); Bruckmueller et al. (2017) |
| Protein amount OATP2B1 [-] | 0.45 | 0.5 | 0.45 | 0.45 | 0.45 | 0.5 | Englund et al. (2006); Drozdzik et al. (2014); Bruckmueller et al. (2017) |
| Transport rate [$\frac{l}{min}$] | 0.067 | 0.19 | 0.19 | 0.012 | 0.012 | - | |

### Model parametrization

Parameter fitting was used to minimize the distance between experimental data and model predictions by optimizing a subset of ten parameters of the model. For this purpose, a sub-dataset of curated time curves of healthy subjects were used which is listed in Tab. 1. Parameter fitting was performed as a two-step procedure based on the route of administration. First, parameters relevant for the intravenous application of talinolol were optimized using the subset of intravenous data. Subsequently, the parameters for oral administration of talinolol were optimized using a subset of talinolol data after oral application.

In the cost function, which is contingent on the parameter p⃗, the sum of the quadratic weighted residuals r*_i,k_* for all time courses k and data points i were minimized. Time courses were weighted by the number of participants in the respective study n*_k_* and individual time points with the standard deviation σ*_i,k_* associated with the measurement resulting in weights w*_i,k_* = n*_k_*/σ*_i,k_*.

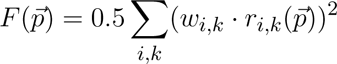

Multiple optimization runs based on a local optimizer were performed for intravenous and oral parameters with the optimal parameters used in the final model. The fitted parameters are provided in Tab. 2.

### Pharmacokinetics parameters

Pharmacokinetic parameters of talinolol were calculated from the plasma-concentration time courses and urinary excretion using standard non-compartmental methods. The elimination rate k_el_[1/min] was calculated via linear regression in logarithmic space in the decay phase. The area under the curve AUC [mmole*·*min/L] was calculated via the trapezoidal rule and extrapolated to infinity based on linear interpolation. Clearance Cl [ml/min] was calculated as Cl = k_el_ *·* V_d_ with the apparent volume of distribution V_d_ = D/(AUC*_∞_ ·* k*_el_*). D is the applied dose of talinolol.

## 3 RESULTS

In this work, an extensive database of talinolol pharmacokinetics was established and used to develop and validate a physiologically based pharmacokinetic (PBPK) model of talinolol.

### 3.1 Database of talinolol pharmacokinetics

Via a dedicated literature search a large number of studies related to the pharmacokinetics of talinolol could be identified. From the initial corpus of 130 studies, duplicates were removed and studies for which the full-text PDF was not accessible were excluded. The remaining 111 studies were screened based on our inclusion criteria, i.e., studies must be performed in human subjects, data must be measured in vivo, and studies must report talinolol time courses. In addition to studies in healthy subjects, studies in patients with renal function impairment or after cholecystectomy were included. In the study reporting pharmacokinetic data in cholecystectomy patients, bile was collected via a T-drain, providing information on the enterohepatic circulation of talinolol Wetterich et al. (1996). One study was excluded because of an incorrect fasting protocol, and another study was excluded because of the administration of radioactive talinolol. This resulted in 33 clinical studies that were curated and formed the database for the development and evaluation of the PKDB model. An overview of the PRISMA flow diagram is provided in Fig. S1. Tab. 1 provides an overview of the curated studies, such as number of subjects, dosing protocol, route of administration, and genetic variants. All data have been uploaded to the pharmacokinetics database PK-DB and are freely accessible via the respective PBPK identifier.

### 3.2 PBPK model of talinolol

Based on the dataset, a physiologically based pharmacokinetic (PBPK) model for talinolol was developed (Fig. 1). The model is hierarchically organized, with the top layer representing the whole body, including lung, liver, kidneys, intestine, and a rest compartment. Transport of talinolol is modeled via the systemic circulation.

Talinolol can be administered in the model either intravenously, orally or by intestinal infusion. After oral administration, the dissolved drug enters the intestine. In the intestinal lumen, talinolol is imported into enterocytes via OATP2B1 located on the apical membrane. Subsequently, a fraction of talinolol is transported back into the intestinal lumen via P-gp efflux transport. After passage through the intestine the unabsorbed fraction is excreted in the feces (Fig. 1B). Intestinal infusion of talinolol involves insertion of a tube into a specific segment of the intestine Gramatté et al. (1996); Bogman et al. (2005). To simulate this procedure, the intestinal model consisted of multiple segments corresponding to different sections of the intestine, i.e. the duodenum, jejunum 1, jejunum 2, ileum 1, ileum 2, and colon. Each segment has specific parameters, transport rates, and varies in length, volume, and enzymatic activity of individual transporters, with parameters listed in Tab. 3.

Talinolol enters the liver either via the portal vein (e.g. after oral administration) or via the hepatic artery. Active uptake of talinolol from plasma into hepatocytes occurs via OATP1B1. Metabolism of talinolol accounts for less than 1 % and recovery of talinolol metabolites in urine is negligible, therefore metabolism of talinolol was not included in the model. Biliary export of talinolol from the liver to the duodenum completes the enterohepatic circulation where talinolol can be reabsorbed or excreted (Fig. 1C). Excretion of talinolol from the bloodstream into the urine is described in the renal model (Fig. 1D).

### 3.3 Model performance

The model allows accurate prediction of talinolol pharmacokinetics following administration of talinolol intravenously (Fig. 2), after single doses orally(Fig. 3), and after multiple doses orally (Fig. 4) ranging from 25 mg to 400 mg. The model predictions include time courses of talinolol in plasma and bile, as well as the amount of talinolol excreted in urine, feces, and bile. The model predictions show very good agreement with clinical data from healthy controls from Bernsdorf et al. (2006); de Mey et al. (1995); Fan et al. (2009b); Giessmann et al. (2004); Han et al. (2009); He et al. (2007, 2012); Krueger et al. (2001); Nguyen et al. (2015); Schwarz et al. (1999, 2000, 2007); Siegmund et al. (2003); Trausch et al. (1995); Tubic et al. (2006b); Westphal et al. (2000a,b); Wetterich et al. (1996); Xiao et al. (2012); Yan et al. (2013); Zhang et al. (2005); Zeng et al. (2009); Zschiesche et al. (2002).

**Figure 2:**
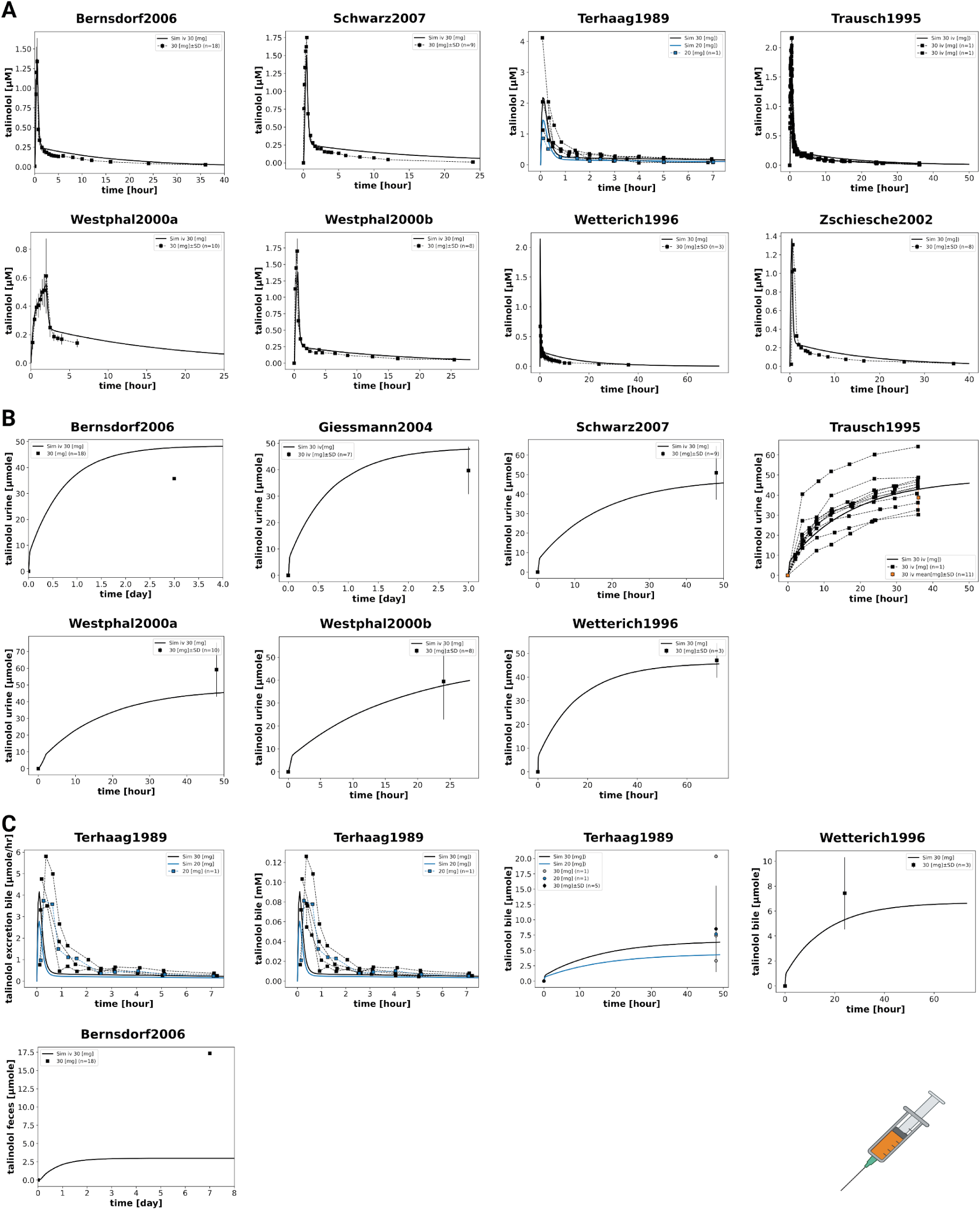
Model predictions of data after intravenous talinolol application. Each subject received a single dose of 30 mg talinolol. One subject received a lower dose of 20 mg (blue) (Terhaag et al., 1989). Model prediction as solid line, data as dashed line. **(A)** talinolol plasma concentration, **(B)** urinary excretion of talinolol, **(C)** fecal and biliary excretion of talinolol. Data from Bernsdorf et al. (2006); Schwarz et al. (2007); Terhaag et al. (1989); Trausch et al. (1995); Westphal et al. (2000a,b); Wetterich et al. (1996); Zschiesche et al. (2002); Giessmann et al. (2004).

**Figure 3:**
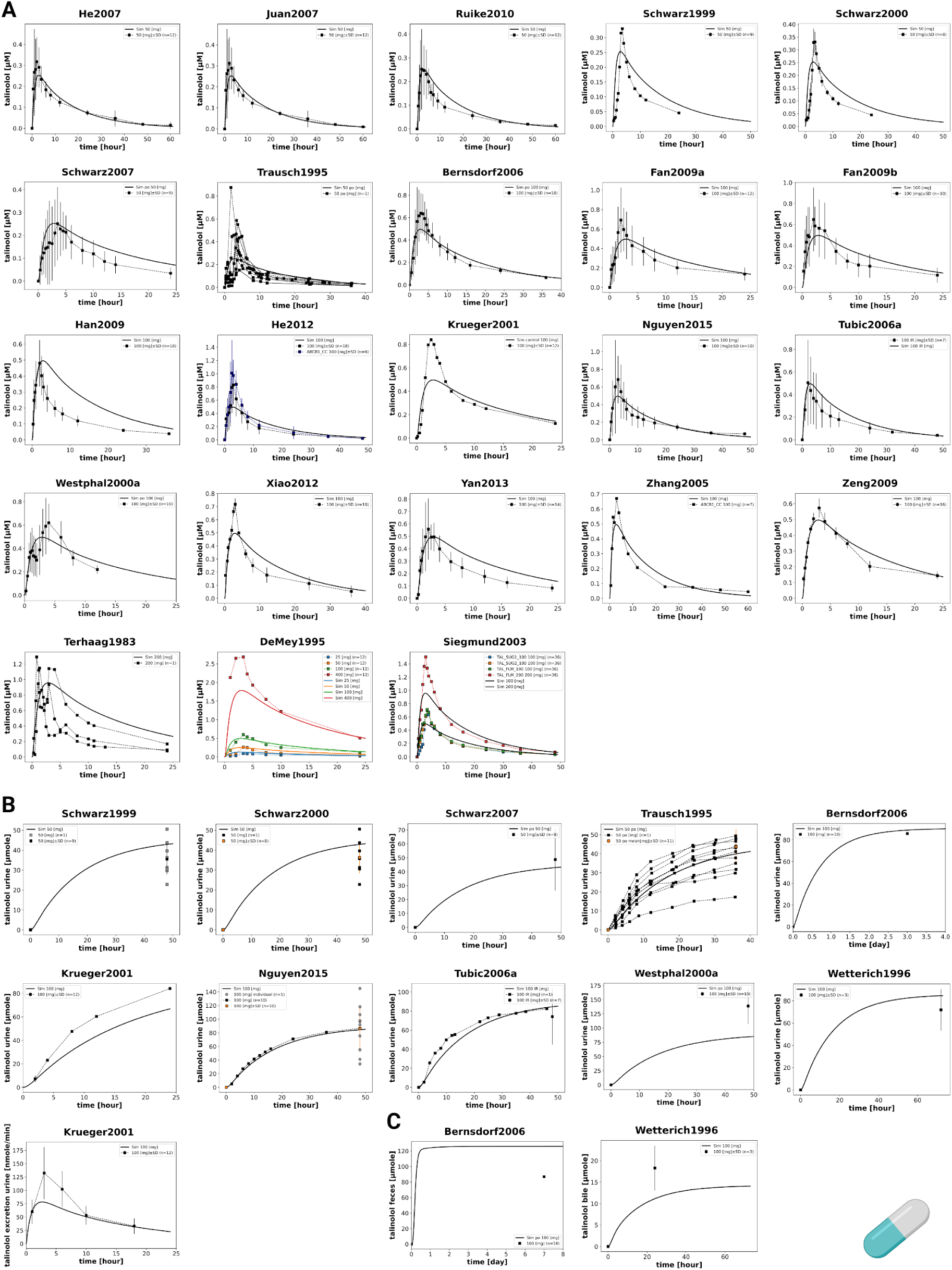
Model predictions of data after oral application of a single dose of talinolol. Subjects received talinolol doses ranging from 25 to 400 mg, with different colors corresponding to different doses. Model prediction as solid line, data as dashed line. **(A)** talinolol plasma concentration, **(B)** urinary excretion, and **(C)** fecal and biliary excretion of talinolol. Data from He et al. (2007); Juan et al. (2007); Ruike et al. (2010); Schwarz et al. (1999, 2000, 2007); Trausch et al. (1995); Bernsdorf et al. (2006); Fan et al. (2009a,b); Han et al. (2009); He et al. (2012); Krueger et al. (2001); Nguyen et al. (2015); Tubic et al. (2006b); Westphal et al. (2000a); Xiao et al. (2012); Yan et al. (2013); Zhang et al. (2005); Zeng et al. (2009); Terhaag et al. (1983); de Mey et al. (1995); Siegmund et al. (2003); Wetterich et al. (1996).

**Figure 4:**
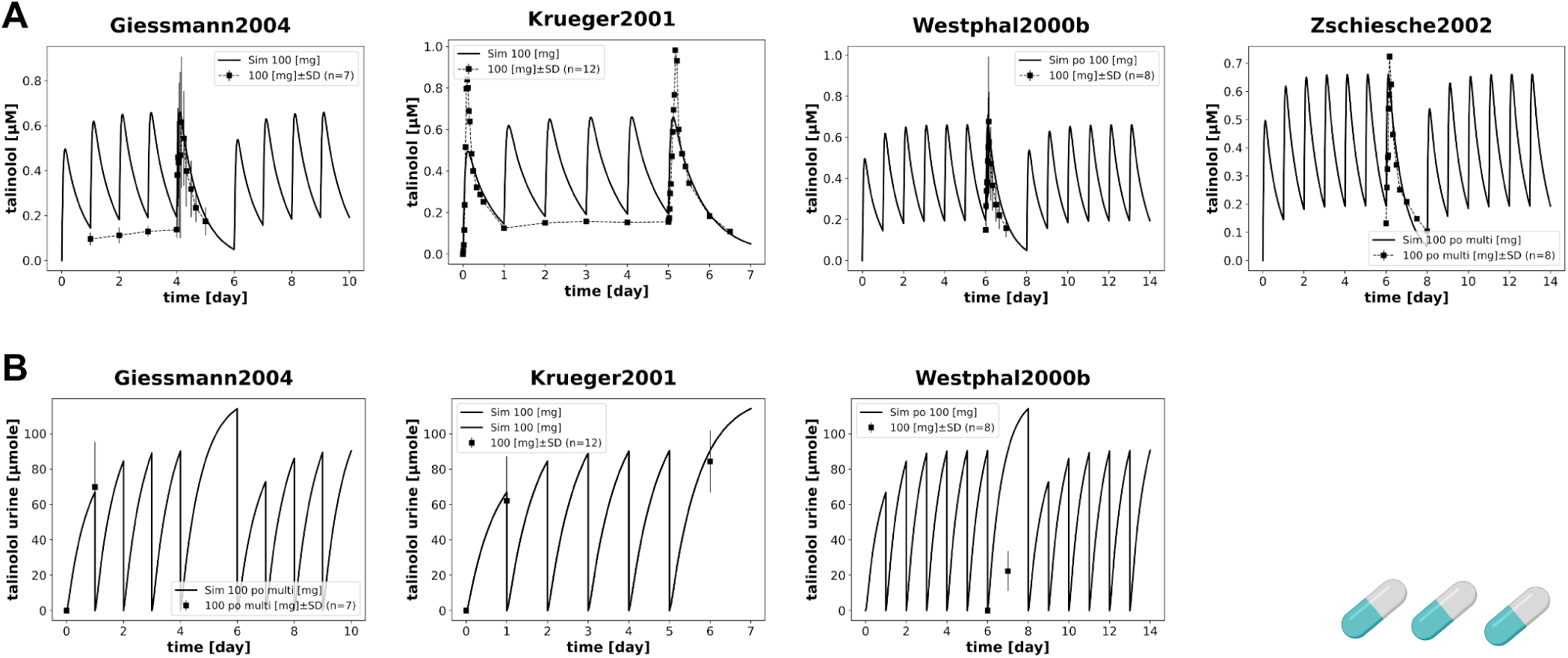
Model prediction of data after oral application of multiple doses of talinolol. Subjects received 100 mg of talinolol daily. Model prediction as solid line, data as dashed line. **(A)** talinolol plasma concentration and **(B)** urinary amounts. Data from Giessmann et al. (2004); Krueger et al. (2001); Westphal et al. (2000b); Zschiesche et al. (2002); Giessmann et al. (2004); Krueger et al. (2001); Westphal et al. (2000b).

Although the renal elimination of talinolol in model is not in perfect agreement with the experimental data from the studies, it is within the range of inter-individual variability observed between the studies. While most of the predictions are in very good agreement with the data, the following discrepancies can be observed: The predicted amount of talinolol excreted in feces is too low after iv application and too high after oral application for Bernsdorf et al. (2006). Second, the plasma concentrations of talinolol are not well predicted for Westphal et al. (2000b), although other multiple dose studies are predicted very well and other data from Westphal et al. (2000b) are in very good agreement with the model predictions.

### 3.4 Effect of P-glycoprotein inhibition and genetic polymorphisms

An important question is how drug-drug interactions with P-gp and genetic polymorphisms of P-gp might affect the pharmacokinetics of talinolol (Fig. 5).

**Figure 5:**
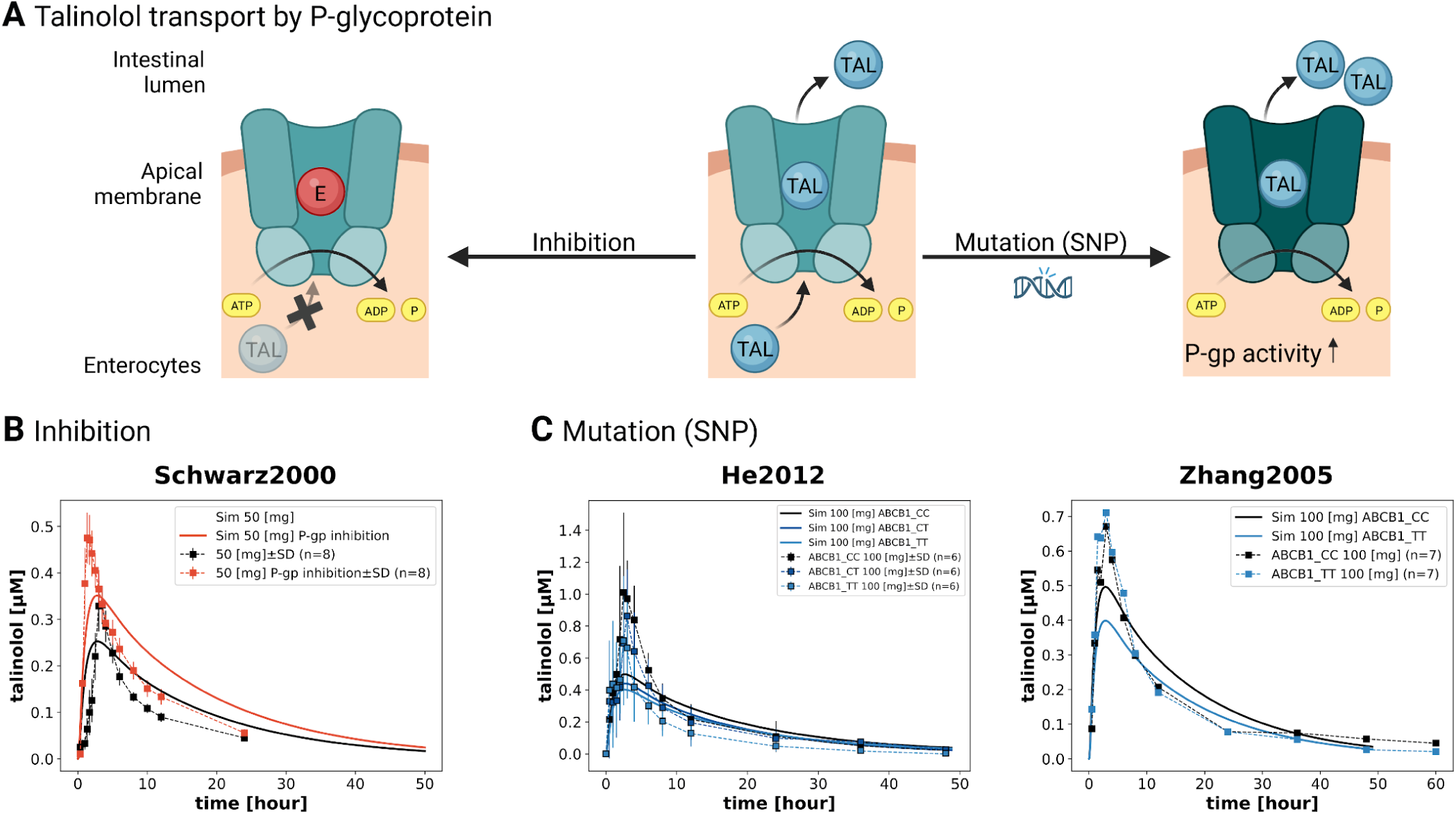
Effect of P-gp inhibition and P-gp genetic polymorphisms on talinolol pharmacokinetics. **(A)** Overview P-gp inhibition and P-gp genetic polymorphisms. Talinolol is transported from enterocytes back into the intestinal lumen by P-gp which can be inhibited by drug-drug interactions such as erythromycin. Alternatively, SNPs may alter P-gp activity. **(B)** Model prediction (solid line) compared to data (dashed line) for P-gp inhibition. The black simulation for normal P-gp activity, in orange for reduced P-gp activity due to inhibition by erythromycin. In the model simulation, P-gp activity was reduced to 50 %. Data from Schwarz et al. (2000). **(C)** Model predictions (solid line) compared to data (dashed line) for P-gp genetic polymorphisms. The analysis was performed on subjects with ABCB1 genotypes 3435CC, 3435CT and 3435TT. For the CT and TT variants, an increased P-gp activity of 20 % and 40 %, respectively, was assumed. Data from (He et al., 2012; Zhang et al., 2005). Created with Biorender.

The inhibition of P-gp by erythromycin was modeled by reducing P-gp activity by 50 % when talinolol was taken orally at a dose of 50 mg. The inhibition of P-gp by co-administration with erythromycin resulted in a higher C_max_ and AUC of talinolol in plasma compared to the administration of talinolol alone. Our model reproduces this effect as shown in Fig. 5B in agreement with the data from Schwarz et al. (2000).

For the genetic polymorphisms of P-gp, we focused on the relationship between ABCB1 genotypes (3435CC, 3435CT, and 3435TT) and P-gp activity. We assumed an increased P-gp activity of 20 % for the CT variant and 40 % for the TT variant (Kim et al. (2001)). The results in Fig. 5C show that an increased number of T alleles correlates with a decrease in AUC. This reduction in plasma concentration for the T variants is in good agreement with the observed reduction in He et al. (2012). In contrast, Zhang et al. (2005) reports no differences between the CC and TT variants.

### 3.5 Effect of P-glycoprotein activity

P-gp activity and cirrhosis were systematically scanned to investigate their effects on the pharmacokinetics of talinolol. The scans were performed with an oral dose of 100 mg of talinolol. Fig. 6B illustrates the effects of different degrees of cirrhosis in relation to P-gp activity on key pharmacokinetic parameters of talinolol, including AUC, k_el_, t_half_, as well as total, renal, and fecal clearance. In the presence of cirrhosis and increased P-gp activity, a marginal decrease in k_el_ and a concomitant increase in t_half_ are observed. While AUC, total clearance and fecal clearance show a dependence on P-gp activity, renal clearance is unaffected by enzymatic activity or the presence of cirrhosis. The higher the P-gp activity, the more talinolol is exported in the intestine via the efflux pump, the lower the absorption and the lower the AUC. As a result of decreased absorption, more talinolol is excreted in the feces.

**Figure 6:**
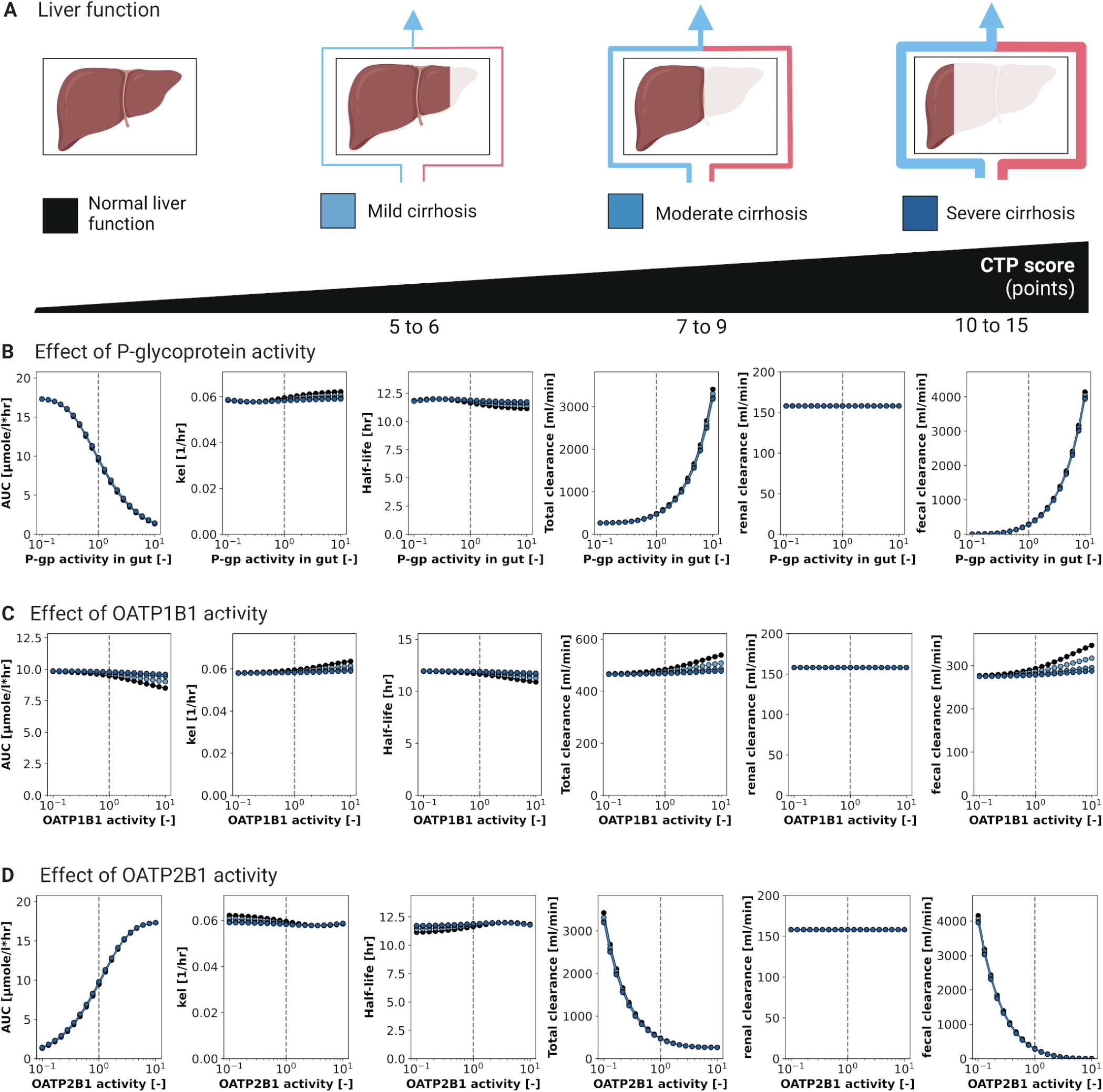
Effect of protein activity on talinolol pharmacokinetics. The effect of changes in transporter activity on the pharmacokinetic parameters AUC, k_el_, t_half_, and total, renal, and fecal clearance was investigated for different degrees of cirrhosis. **(A)** Simulations were performed for normal liver function (black), mild cirrhosis (light blue, CTP A), moderate cirrhosis (medium blue, CTP B), severe cirrhosis (dark blue, CTP C) corresponding to increasing Child-Turcotte-Pugh (CTP) score. **(B)** Effect of P-gp activity. **(C)** Effect of OATP1B1 activity. **(D)** Effect of OATP2B1 activity. Created with Biorender.

An overview of the influence of P-gp activity on the disposition of talinolol in the different compartments after oral or intravenous administration of talinolol is shown in Fig. S2 and Fig. S3.

### 3.6 Effect of OATP1B1 activity

Liver uptake via the OATP1B1 transporter is another important transport mechanism for talinolol in the body. Therefore, our study was designed to investigate the effect of OATP1B1 activity on the pharmacokinetics of talinolol, taking into account the influence of different degrees of cirrhosis at different levels of OATP1B1 activity. The scanning results show that OATP1B1 activity primarily affects talinolol concentration in the liver and the excretion of talinolol into bile (Fig. S4). In particular, induction of OATP1B1 has a strong influence, resulting in increased talinolol concentrations and excretion in liver and bile. In contrast, minimal opposite effects are observed in plasma and urine. The inhibitory activity shows contrasting effects on talinolol disposition in the body. However, talinolol concentrations in individual intestinal segments do not differ significantly based on different levels of OATP1B1 activity. The severity of cirrhosis also has no effect. With increasing severity of cirrhosis, changes are observed only in the liver and the effect of OATP1B1 activity on talinolol disposition in the body is reduced.

An overview of the influence of OATP1B1 activity on the disposition of talinolol in the different compartments after oral or intravenous administration of talinolol is shown in Fig. S4 and Fig. S5.

In the presence of cirrhosis and increased P-gp activity, almost no effect on the pharmacokinetic parameters of talinolol can be observed (Fig. 6C). These findings confirm the previous results and show that with increasing OATP1B1 activity, the severity of cirrhosis plays a role. This leads to a decrease in elimination rate, resulting in lower fecal excretion and prolonged retention of talinolol in the body. Conversely, renal clearance is unaffected by both OATP1B1 activity and the severity of cirrhosis.

### 3.7 Effect of OATP2B1 activity

The last of the three studied transporters is the intestinal uptake transporter OATP2B1. The scanning results show that changes in the enzymatic activity of intestinal OATP2B1 are associated with changes in all the compartments studied. An increase in OATP2B1 activity leads to a marked increase in talinolol absorption, as shown in Supplementary Fig. S6, and consequently to a higher proportion of talinolol being excreted in the urine rather than in the feces. Conversely, a decrease in OATP2B1 activity has the opposite effect, leading to a greater retention of talinolol in the intestinal tract. Notably, this effect becomes more pronounced with increasing distance from the duodenum. The influence of different degrees of cirrhosis on the availability of talinolol in different compartments was also investigated. The scan results show a significant correlation between talinolol concentration and the severity of cirrhosis. Interestingly, talinolol concentration and the influence of OATP2B1 activity tend to decrease with increasing cirrhosis severity.

An overview of the influence of OATP2B1 activity on the distribution of talinolol in different compartments after oral and intravenous administration is shown in Fig. S6 and S7.

Fig. 6D illustrates the effect of cirrhosis severity on pharmacokinetic parameters as a function of OATP2B1 activity. The results show that an increase in OATP2B1 activity leads to an increase in the AUC of talinolol. The influence of reduced OATP2B1 activity on the elimination rate constant (k_el_) and half-life (t_half_) of talinolol in the body depends on the severity of cirrhosis. Under healthy conditions, OATP2B1 activity has minimal effect on k_el_ and t_half_. However, as the severity of cirrhosis increases, k_el_ decreases while t_half_ increases with decreased OATP2B1 activity. Increased OATP2B1 activity is associated with decreased total and fecal clearance of talinolol. Neither OATP2B1 activity nor cirrhosis has a significant effect on renal clearance.

As an important side note, the effects of changes in OATP2B1 activity are opposite to changes in P-gp activity. As expected, an increase in the intestinal influx transporter OATP2B1 has a similar effect as a decrease in the efflux transporter P-gp and vice versa.

### 3.8 Effect of renal function

An important question is how renal function affects the pharmacokinetics of talinolol (Fig. 7). Renal dysfunction is common in patients with cirrhosis. The influence of liver cirrhosis severity in relation to renal function on the pharmacokinetic parameters of talinolol is shown in Fig. 7B. The results indicate that cirrhosis has no additional effect on talinolol pharmacokinetics beyond changes in renal function.

**Figure 7:**
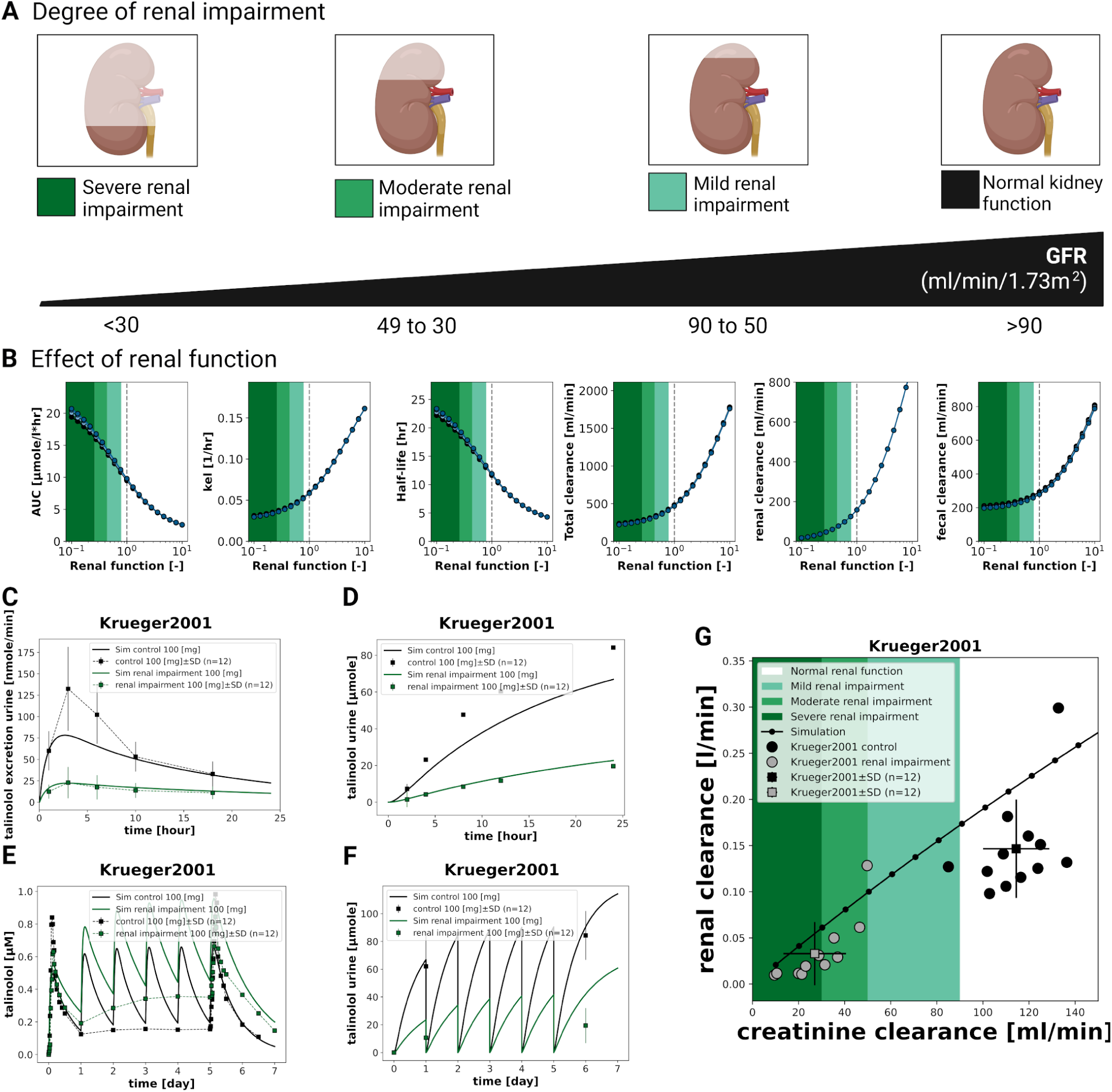
Effect of renal function on talinolol pharmacokinetics. **(A)** Simulations were performed for normal renal function (black), mild renal impairment (light green), moderate renal impairment (medium green), and severe renal impairment (dark green), corresponding to decreasing glomerular filtration rate (GFR). **(B)** The effect of renal function on the pharmacokinetic parameters AUC, k_el_, t_half_ and total, renal and fecal clearance was investigated for different degrees of cirrhosis. Simulations were performed for normal liver function (black), mild cirrhosis (light blue, CTP A), moderate cirrhosis (medium blue, CTP B), severe cirrhosis (dark blue, CTP C) corresponding to increasing Child-Turcotte-Pugh (CTP) score. **(C)** Talinolol urinary excretion rate and **(D)** urine volume after a single dose of talinolol. **(E)** Talinolol plasma concentrations and **(F)** urinary amounts after repeated doses of talinolol. Model predictions as solid lines and data as dashed lines. **(G)** Dependence of the renal clearance of talinolol on the creatinine clearance. Individual subjects as dots and group mean±SD as squares and error bars. The solid black line represents the prediction of the model. Data from Krueger et al. (2001). Created with Biorender.

Renal clearance, on the other hand, depends solely on renal function. As renal function increases, k_el_ and total, renal, and fecal clearance of talinolol also increase, while AUC and t_half_ decrease.

An overview of the influence of renal function on the distribution of talinolol in different compartments after oral and intravenous administration is given in Fig. S8 and Fig. S9.

The model predictions for renal function impairment were validated with experimental data. Krueger et al. (2001) studied the pharmacokinetics of talinolol in 24 subjects with different creatinine clearance values corresponding to different degrees of renal impairment. In Fig. 7C, D, E, F the experimental data with single and repeated administration of talinolol from plasma and urine were compared with the model prediction. Irrespective of the route of administration, it can be seen that with decreasing renal function, the c_max_ in plasma increases and the amount excreted in urine decreases. Overall, the model prediction is in very good agreement with the data. However, it is important to note that the model predicts a slightly lower trajectory but greater accumulation of talinolol in both plasma and urine compared to the data.

Fig. 7G illustrates the relationship between renal clearance and creatinine clearance, which serves as a measure of renal function. In healthy individuals and those with severely impaired renal function, the average renal clearance is approximately 0.03 and 0.15 *<u>^l^</u>*, respectively.

As creatinine clearance improves, the renal clearance of talinolol increases. The model prediction of the dependence of talinolol renal clearance on creatinine clearance (estimated GFR) is in very good agreement with the data.

### 3.9 Site-dependency of intestinal absorption

In addition to oral administration of talinolol in solid form, it is possible to infuse talinolol specifically into different intestinal segments. The intestinal site dependency scan as a function of P-gp activity shown in Fig. 8 allows a detailed investigation of the pharmacokinetics of talinolol after administration into the individual segments, duodenum, jejunum 1, jejunum 2, ileum 1, ileum 2 and colon. The infusion is given at a constant rate for five hours. In the study of site-specific infusions, an immediate decrease in talinolol concentration is observed in plasma, liver, bile, and individual intestinal segments after the end of the infusion, except in urine and feces, which continue to increase after the end of the infusion due to the cumulative nature of the amounts (i.e., total amount in feces and urine). In addition, minimal talinolol was detected in the regions prior to the infusion site due to the minimal enterohepatic circulation of talinolol.

**Figure 8:**
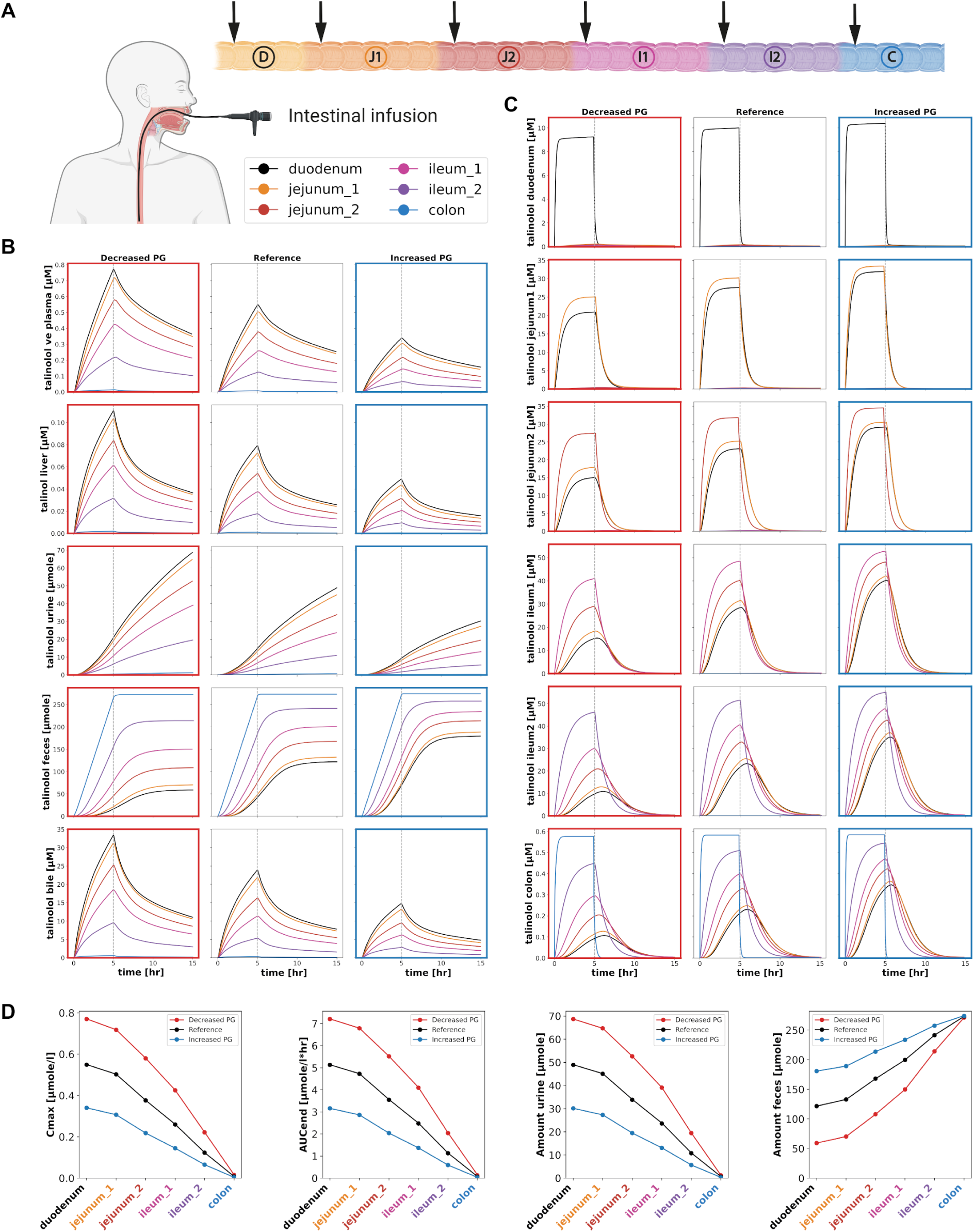
Effect of site-specific intestinal infusion on talinolol pharmacokinetics. **(A)** Intestinal infusion is administered through a tube placed orally at a specific site along the gastrointestinal tract. A 100 mg infusion of talinolol was simulated over a five-hour period. Site-specific intestinal infusion in the duodenum (black), jejunum 1 (orange), jejunum 2 (red), ileum 1 (pink), ileum 2 (purple), and colon (blue) are shown. Simulations were performed for decreased P-gp activity (0.5, red box), reference P-gp activity (1.0, black box), and increased P-gp activity (2.0, blue box). **(B)** Talinolol in venous plasma, liver, urine, feces, and bile. **(C)** Talinolol concentration in each intestinal segment. **(D)** The effect of site-specific infusion on the pharmacokinetic parameters C_max_, AUC, and amount in urine and feces depending on the localization of the intestinal infusion and P-gp activity. Created with Biorender.

Overall, the results (Fig. 8B) show a decrease in talinolol concentration in plasma, urine and bile with increasing distance of the infusion site from the duodenum. In contrast, the amount of talinolol in the feces increases. This observation is further supported by the infusion of talinolol into the colon, where no talinolol is absorbed into the body and 100 % of the administered talinolol is recovered in the feces. While fecal excretion remains constant during colonic infusion, the bioavailability of talinolol decreases both with distance from the infusion site to the duodenum and with increasing P-gp activity.

The site of infusion in the intestine and P-gp activity have strong effects on the pharmacokinetic parameters of talinolol.

With increasing distance of the infusion site from the duodenum, the c_max_, AUC, and amount in urine decrease, while the amount in feces increases. A decrease in P-gp activity increases c_max_, AUC, and urine amount due to additional absorption when the efflux transporter activity is decreased.

To validate the site-specific model, data from intestinal infusion studies were used (Fig. 9). In addition to studies in healthy subjects, studies in patients with renal impairment or cholecystectomy were included. In the study reporting pharmacokinetic data in cholecystectomy patients, bile was collected via a T-drain, providing information on the enterohepatic circulation of talinolol Wetterich et al. (1996). Gramatté et al. (1996) infused either in the duodenum or in the upper part of the jejunum, whereas Bogman et al. (2005) infused exclusively in the duodenum. Significant differences in serum talinolol concentrations were observed among subjects. In all cases, less talinolol was absorbed when the infusion was initiated distal to the duodenum. Although the model predictions were not in good agreement with the absolute plasma concentrations, the relative decrease in talinolol plasma concentrations when the infusion site was varied can be recapitulated by the model, see Fig. 9B, D.

**Figure 9:**
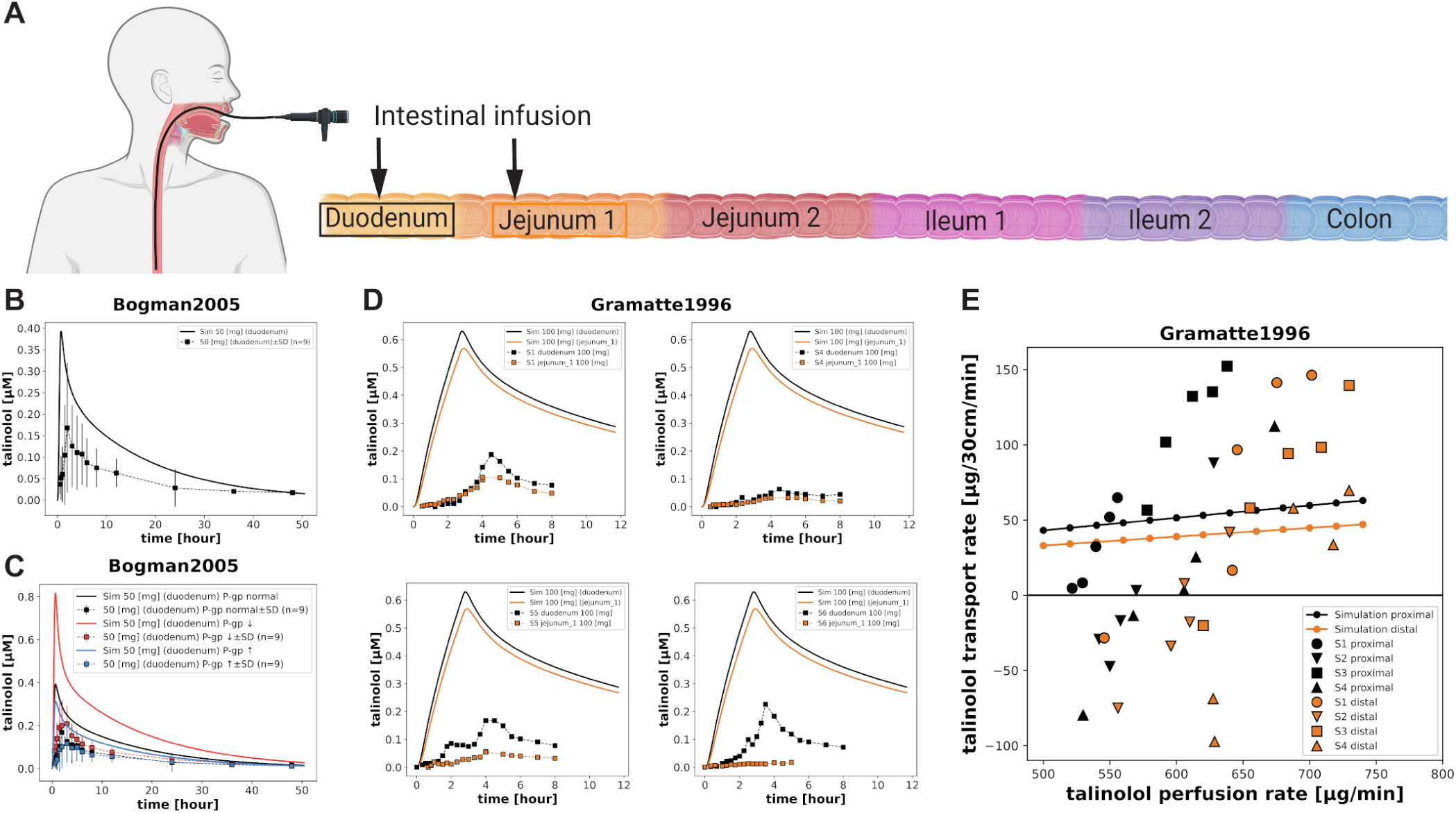
Site-specific intestinal infusions. **(A)** For model validation, site-specific infusions of talinolol were simulated in the duodenum (black, proximal) or upper jejunum (orange, distal). **(B)** Administration of 50 mg talinolol over 25 minutes by intestinal infusion in the duodenum (black) according to the protocol of Bogman et al. (2005). **(C)** Effect of reduced (red) and increased (blue) P-gp activity due to coadministration of 0.04 % TPGS and 0.08 % poloxamer 188, respectively. In the simulation, the activity was reduced to 0.1915 and increased to 1.4 in accordance with the reported in vitro data. **(D)** Administration of 100 mg talinolol over 160 minutes by intestinal infusion in four subjects in either the duodenum (black) or upper jejunum (orange) according to the protocol of Gramatté et al. (1996). **(E)** Transepithelial transport of talinolol as a function of talinolol perfusion rate and localization within the 30 cm test segment according to the protocol of Gramatté et al. (1996). Data from Bogman et al. (2005); Gramatté et al. (1996). Created with Biorender.

Figure 9C illustrates the modulation of P-gp activity, showing a 40 % increase with 0.08 % poloxamer 188 co-administration and a 19.15 % decrease with 0.04 % TPGS co-administration, as reported in the *in-vitro* of Bogman et al. (2005). Examination of the relationship between the net absorption rate of talinolol and the perfusion rate and location, shown in Fig. 9E, reveals an increased transport rate in the proximal compared to the distal region and an increase in transport rate with increasing perfusion rate, consistent with the data. Finally, we examined the effect of intestinal perfusion rate on the net talinolol transport rate in the intestine. The predicted net transport rates fall within the scatter of the data point cloud from Gramatté et al. (1996), although negative transport rates and a much larger slope are present in the data. The qualitative effect of and increase in transport rate with perfusion rate and a shift to a lower transport rate from proximal to distal infusion is shown by the model. These site-selective differences in uptake were effectively described by the model.

## 4 DISCUSSION

In this work, an extensive quantitative data set on talinolol was generated and used to develop a PBPK model of talinolol. The data set, which includes 33 studies with a total of 445 subjects, contains information on the time courses and pharmacokinetics following single and multiple oral, intravenous, and intraintestinal administrations of talinolol. To our knowledge, this is the largest available resource on the pharmacokinetics of talinolol with all data freely available from PK-DB. We anticipate that the data set will be an important asset in further studies of talinolol.

Notably, the data have some limitations: (i) Most of the studies were performed in healthy adult subjects, one of the exceptions being a group of patients with renal impairment Krueger et al. (2001). (ii) The concentration-time curves of talinolol in plasma and serum from most studies are consistent with a few exceptions Wang et al. (2013); Schwarz et al. (2005). However, the large standard deviation (when reported) indicates significant inter-individual variability. The same applies to the excretion of talinolol in urine and bile. (iii) Very limited data were available on talinolol recovery in feces, with only Bernsdorf et al. (2006) provided data on the amount of talinolol excreted in feces after both intravenous and oral administration. (iv) Although numerous studies have investigated the pharmacokinetics of talinolol in relation to genetic variants of P-gp, the available data on the time courses of talinolol in relation to P-gp genetic variants are limited. In addition, there is a lack of available data on the relationship between OATP2B1, OATP1B1 and the pharmacokinetic parameters of talinolol. (v) Information on the pharmacokinetics of talinolol in liver disease was very limited and no data were available in cirrhotic patients.

Importantly, our model predictions provide information about possible alterations of pharmacokinetics in the cases were only minimal data is currently available, such as changes in OATP2B1 activity, P-gp activity, or hepatic impairment. Our simulations could therefore fill an important gap of knowledge in talinolol pharmacokinetics and motivate target experiments. Hopefully, future research will address these areas thereby allowing a more comprehensive validation of the model and understanding of talinolol.

A physiologically based pharmacokinetic (PBPK) model of talinolol was developed based on the established data set. The model was used to investigate the influence of several factors on talinolol pharmacokinetics: (i) P-gp genetic variants; (ii) inhibition of P-gp; (iii) activity of OATP2B1 and OATP1B1; (iv) effect of disease, namely hepatorenal impairment; and (v) site-specific distribution of P-gp and OATP2B1 in the intestine. The model accurately predicts the concentration-time profile of talinolol after oral and intravenous administration and after single and multiple dosing. Furthermore, the model accurately describes the effect of genetic variants of P-gp on the pharmacokinetics of talinolol, the effect of inhibition of P-gp, the effect of renal impairment, as well as site-specific infusion of talinolol in the intestine.

The model was fitted using single-dose intravenous and oral data only from immediate-release tablets. As a result, the model is not applicable to sustained-release formulations, which have significantly different pharmacokinetics with later and lower peak plasma concentrations of talinolol. The model could easily be extended to include such data, e.g. by adjusting the dissolution and absorption rates of the tablets based on the tablet formulation. Because only a small subset of data was reported for slow-release tablets, the model focused on the immediate-release formulations.

The model presented is an average model and did not account for intra-individual variability, despite the large variability observed in the curated data. Future model extensions could account for this variability, e.g. using nonlinear mixed effects models or sampling from underlying parameter distributions.

An important part of the model was the subdivision of the intestine into subcompartments corresponding to the duodenum, jejunum, ileum and colon, which allowed to model the effect of site-specific intestinal infusion of talinolol and the effect of site-dependent influx and efflux transporters. The protein level of OATP2B1 remains relatively constant in the different intestinal segments, whereas the protein level of P-gp increases along the small intestine. This distribution of OATP2B1 as an importer and P-gp as an efflux transporter in the intestine has the consequence that the bioavailability of talinolol in plasma decreases with increasing distance from the duodenum. The model successfully described these differences. Such site-specific absorption windows may be important determinants of drug pharmacokinetics. Interestingly, based on the site dependency scan (see Fig. 8), the model shows that talinolol appears in very small amounts in segments above the infusion site due to the unidirectional motion within the intestinal model combined with minimal enterohepatic circulation. This finding suggests that enterohepatic circulation is relatively insignificant in the disposition of talinolol.

It is important to note that the intestinal model only includes the two primary transporters P-gp and OATP2B1, which are the main transporters reported to be involved in the intestinal disposition of talinolol. The possible role of alternative transporters, such as OATP1A2 Shirasaka et al. (2010) or MRP2 Giessmann et al. (2004), was not considered in the model.

The present study focused on investigating the influence of changes in activity of P-gp, OATP2B1 and OATP1B1 on the pharmacokinetics of talinolol, e.g. due to genetic polymorphisms. The results concerning the P-gp genotype were particularly interesting. However, conflicting results have been reported regarding the effect of P-gp variants. Kim et al. (2001) showed that the mutant genotype 1236 C>T, 2677 G>T, 3435 C>T (TTT) showed a 40 % reduction in AUC compared to the wild type genotype (CCC). This mutation-induced change was modeled and compared with the studies of He et al. (2012) and Zhang et al. (2005). The model showed good agreement with the data from He et al. (2012). It is worth noting that He et al. (2012) only examined the genetic variants of the SNP at position 3435. The effect of OATP1B1 gene polymorphisms on talinolol pharmacokinetics has only been reported in a single study Bernsdorf et al. (2006). The model successfully confirmed the results by showing a correlation between the predicted half-life of talinolol and the activity of OATP1B1, as shown in Fig. 6B. Unfortunately, the data were very limited and no time course data were reported that would have allowed a more thorough validation of our predictions. Given the limited sample size, future investigations should consider further exploring the effects of genetic variants of OATP1B1 on the pharmacokinetics of talinolol.

An important question of this study was how hepato-renal impairment affects the pharmacokinetics of talinolol. Our model suggests that a higher degree of hepatic impairment results in a decreased uptake of talinolol into the liver. This could result in a prolonged drug residence time in the body and higher plasma levels.

The model shows a clear correlation between renal clearance of talinolol and renal function. Furthermore, renal function exerts a particularly strong influence on the AUC of talinolol because the main route of elimination of talinolol is through the kidneys. When compared (see Fig. 7) with Krueger et al. (2001), who collected an extensive pharmacokinetic data set of talinolol in subjects with varying renal function, the difference in AUC after single and multiple dosing of talinolol was accurately captured for different renal functions. In even better agreement, the model prediction is consistent with the observed data regarding the dependence of renal clearance on creatinine clearance, a well-established indicator of renal function.

The present work provides valuable insights into the influence of cirrhosis and renal function on the pharmacokinetics of talinolol, particularly in relation to the activity of P-gp, OATP2B1, and OATP1B1. The results highlight the need for further research in this area to optimize the use of talinolol in patients with cirrhosis and renal insufficiency and to ensure effective therapy. Our results provide important insights into factors affecting talinolol pharmacokinetics and talinolol-based metabolic phenotyping.

## AUTHOR CONTRIBUTIONS

BSM and MK conceived and designed the study, developed the computational model, curated the data, implemented and performed the analysis, and drafted the manuscript. JG provided support with PK-DB, data curation, and modeling. All authors actively participated in the discussions of the results, contributed to critical revisions of the manuscript, and approved the final version for submission.

## DATA AVAILABILITY STATEMENT

The datasets analyzed for this study can be found in PK-DB available from https://pk-db.com.

## Supplementary Material

### 1 SUPPLEMENTARY FIGURES

**Figure S1.**
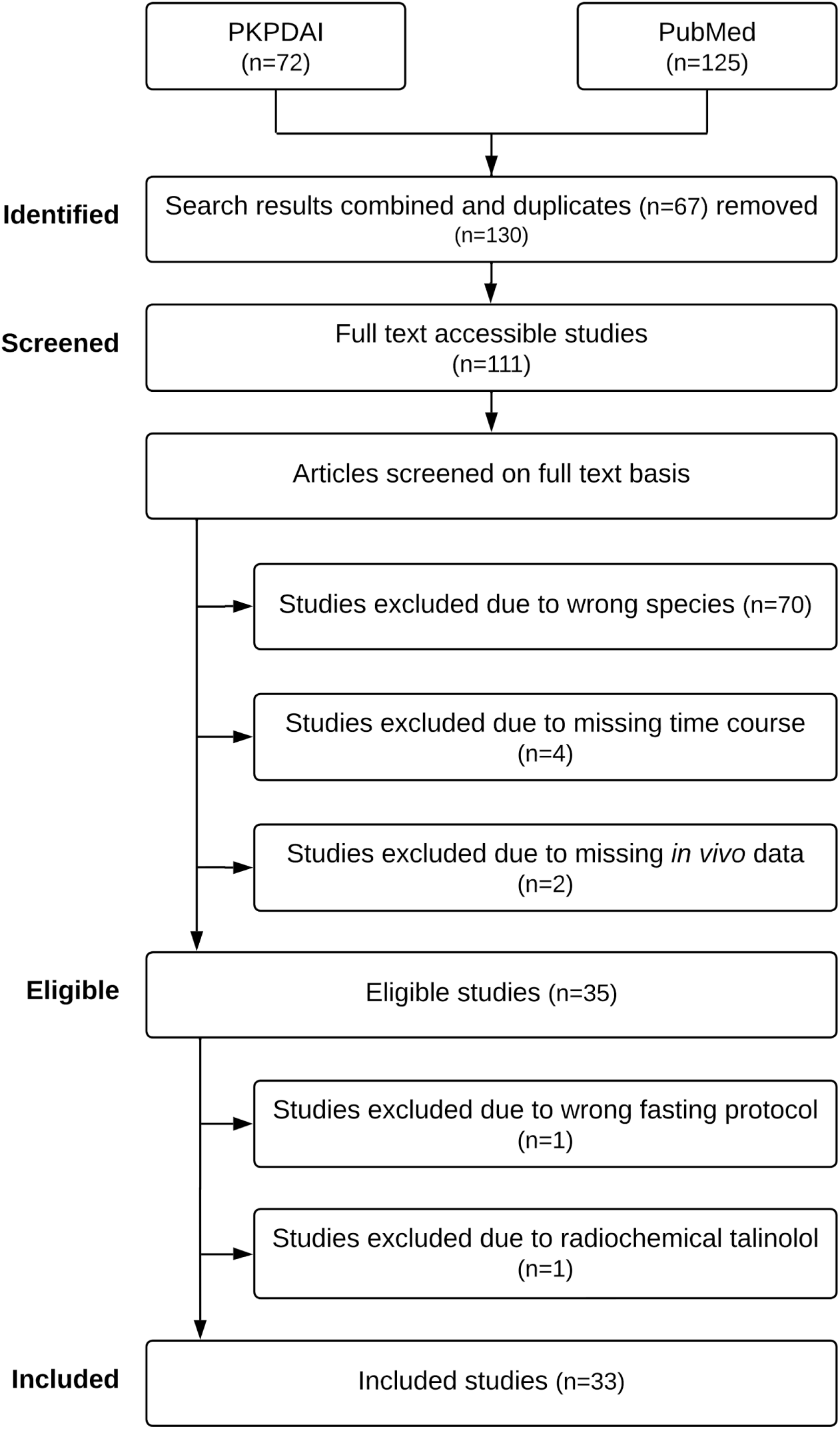
PRISMA flowchart. Overview of the literature search and data selection for the established talinolol pharmacokinetic dataset. PubMed (search term `pharmacokinetics AND talinolol`) and PKPDAI (search term `talinolol`) were used for the initial literature search. Application of the eligibility criteria resulted in 35 studies, of which 33 were curated.

**Figure S2.**
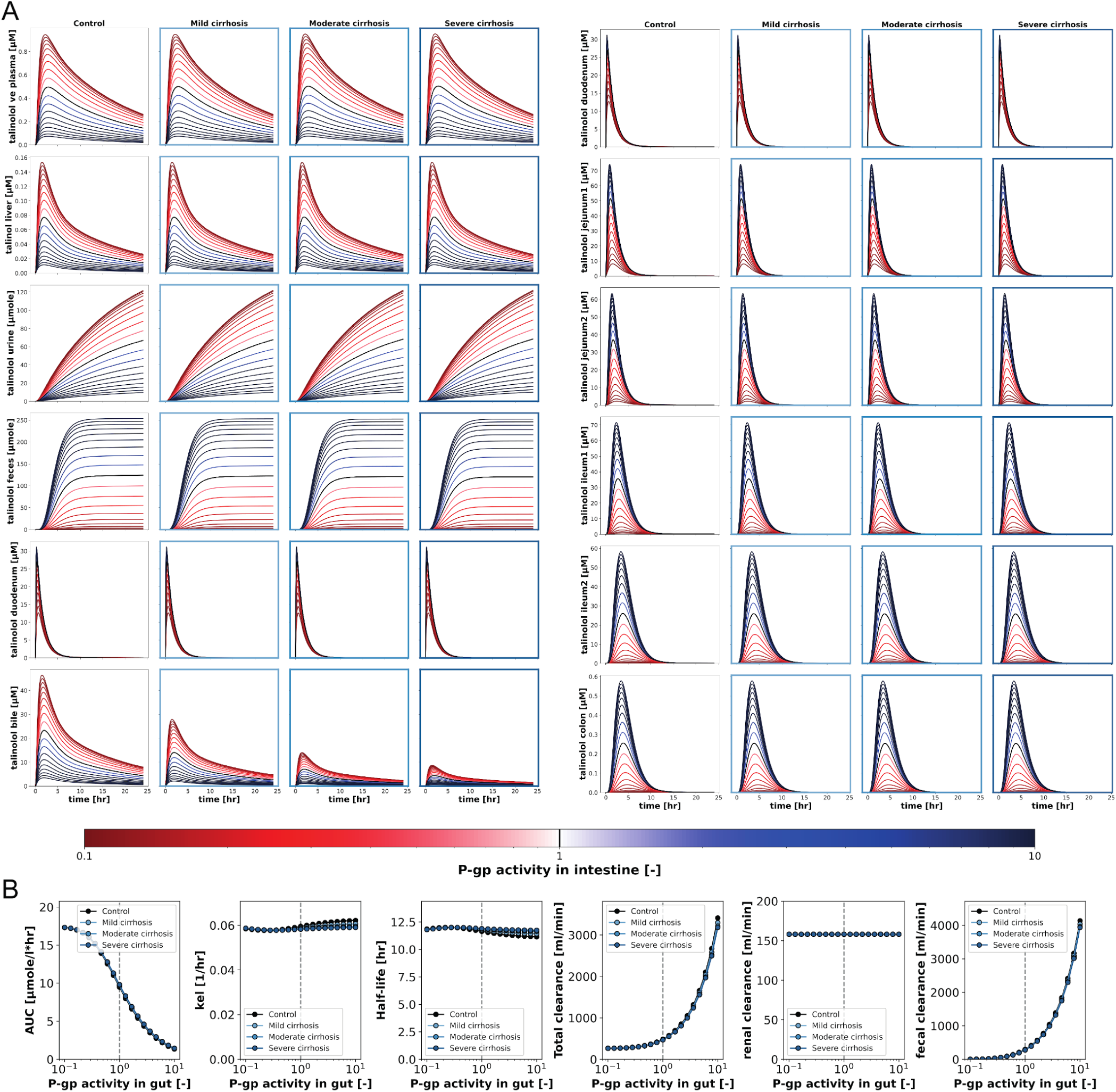
Scan of P-glycoprotein activity after oral administration of talinolol. 100 mg of talinolol was administered orally and P-gp activity was varied as `np.logspace(start=-1, end=10, num=19)` corresponding to a scan of `GU f PG` from 0.1 to 10. **(A)** Overview of the effects of protein activity on the time course of talinolol in different compartments at different degrees of cirrhosis. Increased and decreased protein activities are indicated by the blue and red gradients, respectively. **(B)** Overview of the effects of protein activity on talinolol pharmacokinetic parameters in different degrees of cirrhosis: Area under the curve (AUC), elimination rate (k_el_), half-life (t_half_) and clearance (renal, faecal and total).

**Figure S3.**
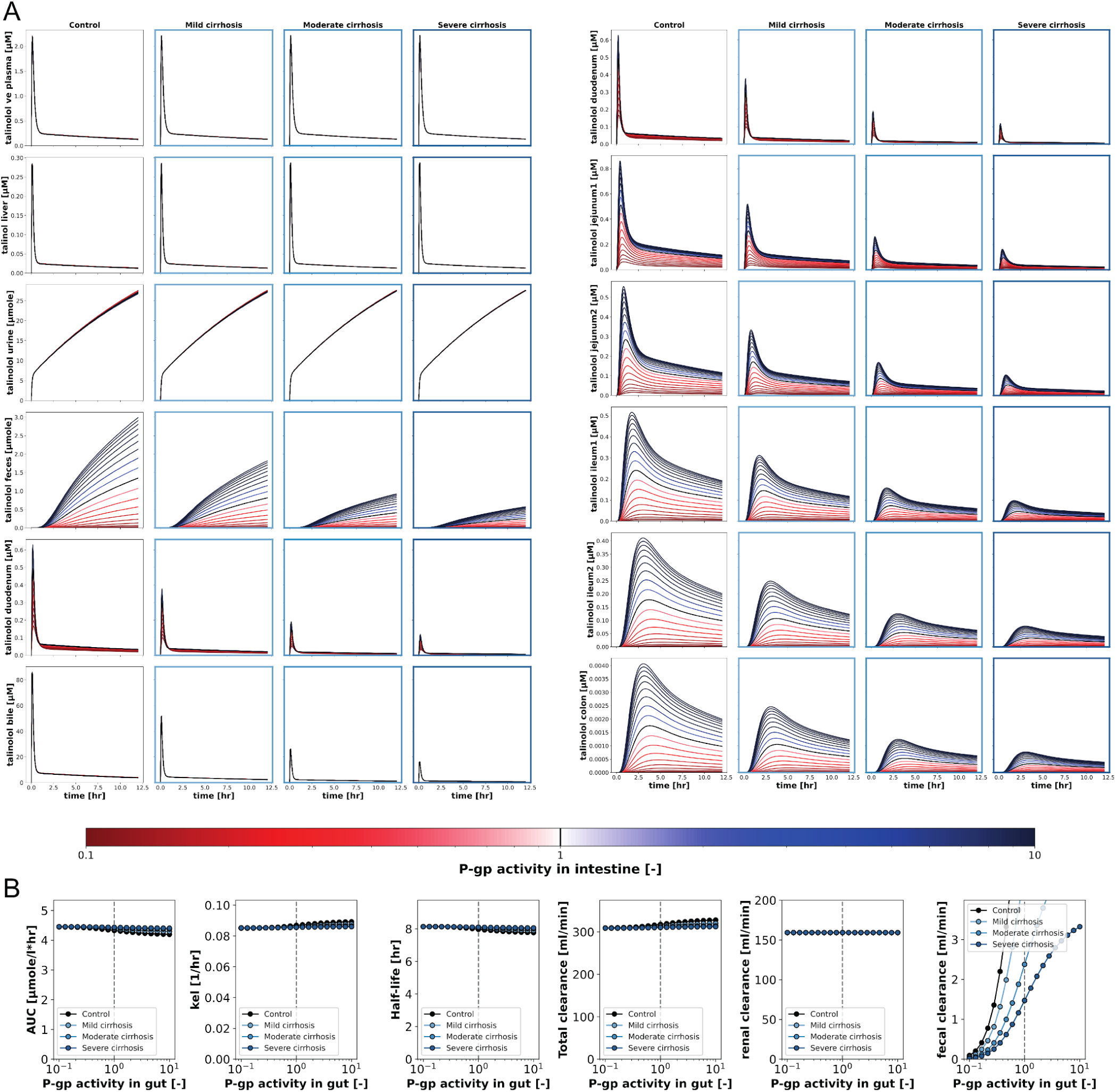
Scan of P-glycoprotein activity after intravenous administration of talinolol. 30 mg of talinolol was administered intravenously and P-gp activity was varied as `np.logspace(start=-1, end=10, num=19)` corresponding to a scan of `GU f PG` from 0.1 to 10. **(A)** Overview of the effects of protein activity on the time course of talinolol in different compartments at different degrees of cirrhosis. Increased and decreased protein activities are indicated by the blue and red gradients, respectively. **(B)** Overview of the effects of protein activity on talinolol pharmacokinetic parameters in different degrees of cirrhosis: Area under the curve (AUC), elimination rate (k_el_), half-life (t_half_) and clearance (renal, faecal and total).

**Figure S4.**
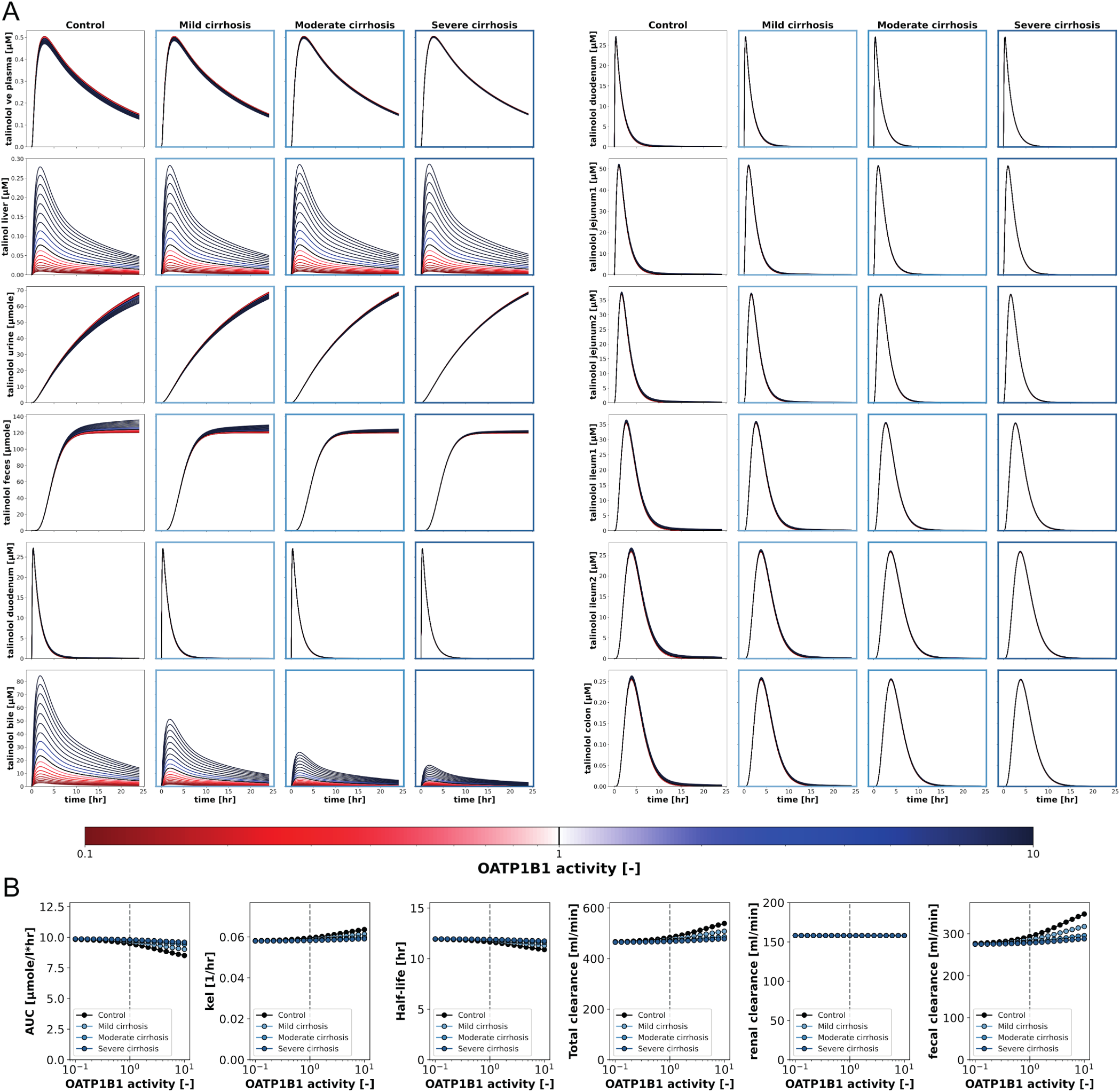
Scan of OATP1B1 activity after oral administration of talinolol. 100 mg of talinolol was administered orally and OATP1B1 activity was varied as `np.logspace(start=-1, end=10, num=19)` corresponding to a scan of `LI f OATP1B1` from 0.1 to 10. **(A)** Overview of the effects of protein activity on the time course of talinolol in different compartments at different degrees of cirrhosis. Increased and decreased protein activities are indicated by the blue and red gradients, respectively. **(B)** Overview of the effects of protein activity on talinolol pharmacokinetic parameters in different degrees of cirrhosis: Area under the curve (AUC), elimination rate (k_el_), half-life (t_half_) and clearance (renal, faecal and total).

**Figure S5.**
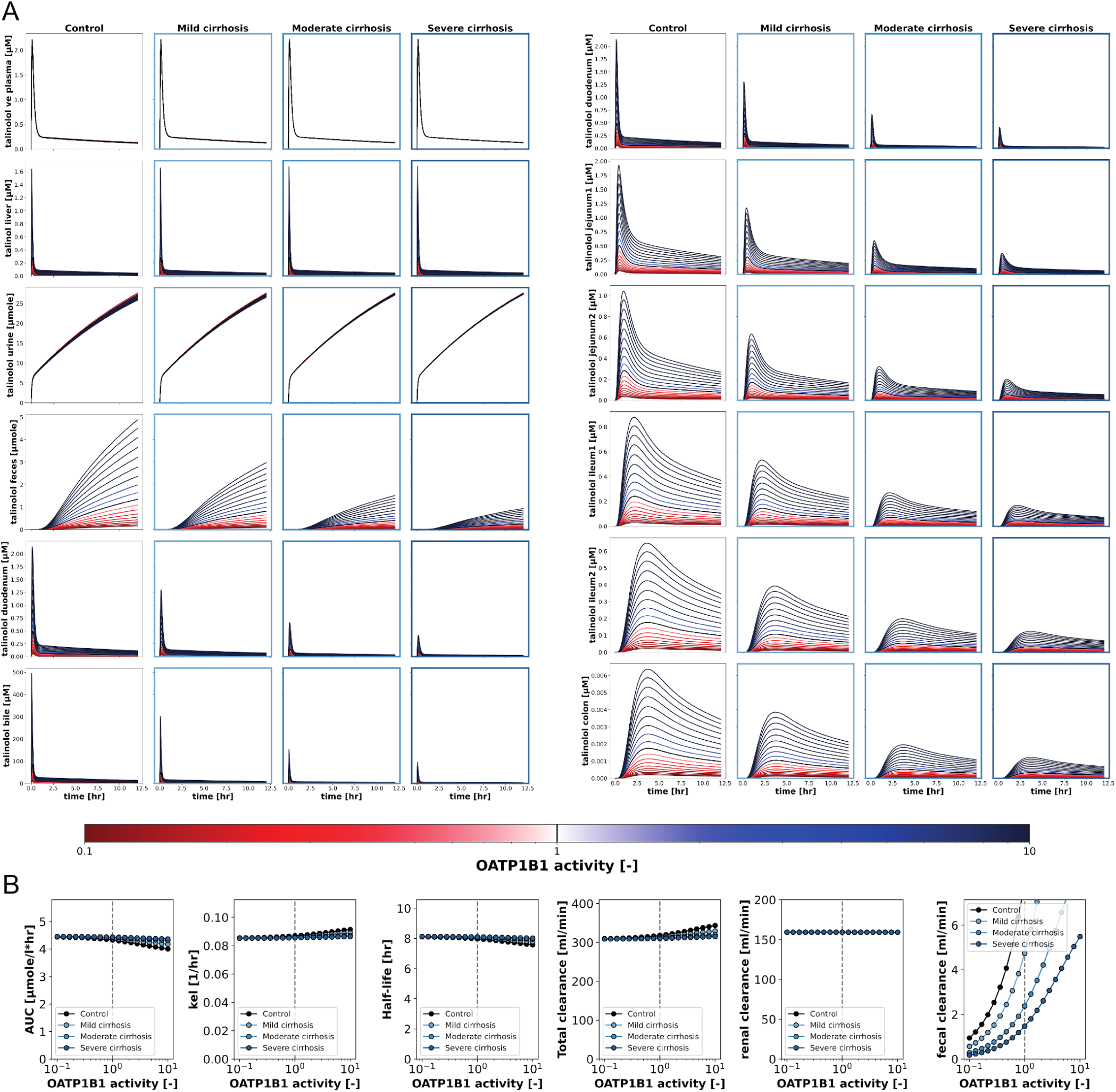
Scan of OATP1B1 activity after intravenous administration of talinolol. 30 mg of talinolol was administered intravenously and OATP1B1 activity was varied as `np.logspace(start=-1, end=10, num=19)` corresponding to a scan of `LI f OATP1B1` from 0.1 to 10. **(A)** Overview of the effects of protein activity on the time course of talinolol in different compartments at different degrees of cirrhosis. Increased and decreased protein activities are indicated by the blue and red gradients, respectively. **(B)** Overview of the effects of protein activity on talinolol pharmacokinetic parameters in different degrees of cirrhosis: Area under the curve (AUC), elimination rate (k_el_), half-life (t_half_) and clearance (renal, faecal and total).

**Figure S6.**
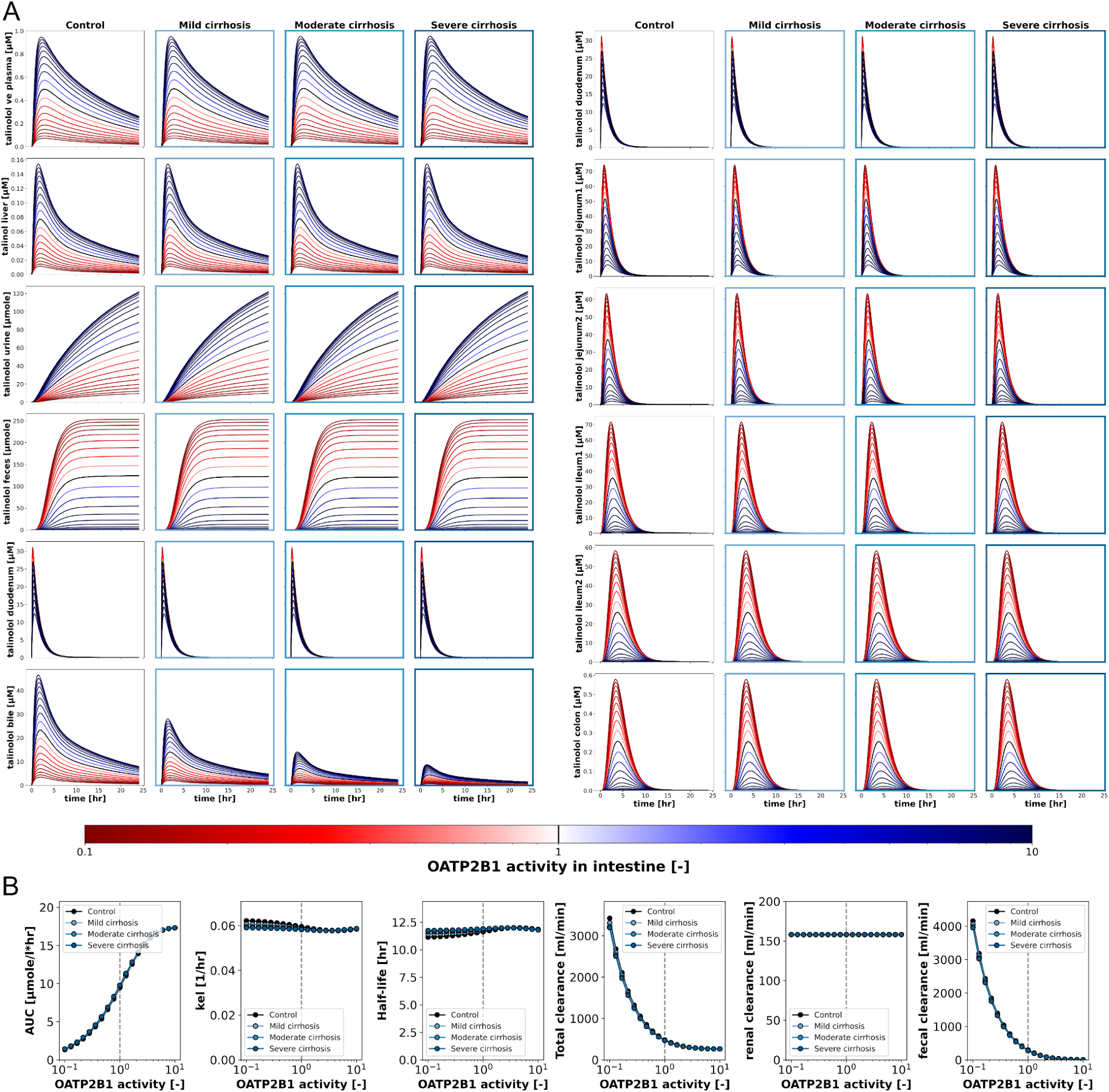
Scan of OATP2B1 activity after oral administration of talinolol. 100 mg of talinolol was administered orally and OATP2B1 activity was varied as `np.logspace(start=-1, end=10, num=19)` corresponding to a scan of `GU f OATP2B1` from 0.1 to 10. **(A)** Overview of the effects of protein activity on the time course of talinolol in different compartments at different degrees of cirrhosis. Increased and decreased protein activities are indicated by the blue and red gradients, respectively. **(B)** Overview of the effects of protein activity on talinolol pharmacokinetic parameters in different degrees of cirrhosis: Area under the curve (AUC), elimination rate (k_el_), half-life (t_half_) and clearance (renal, faecal and total).

**Figure S7.**
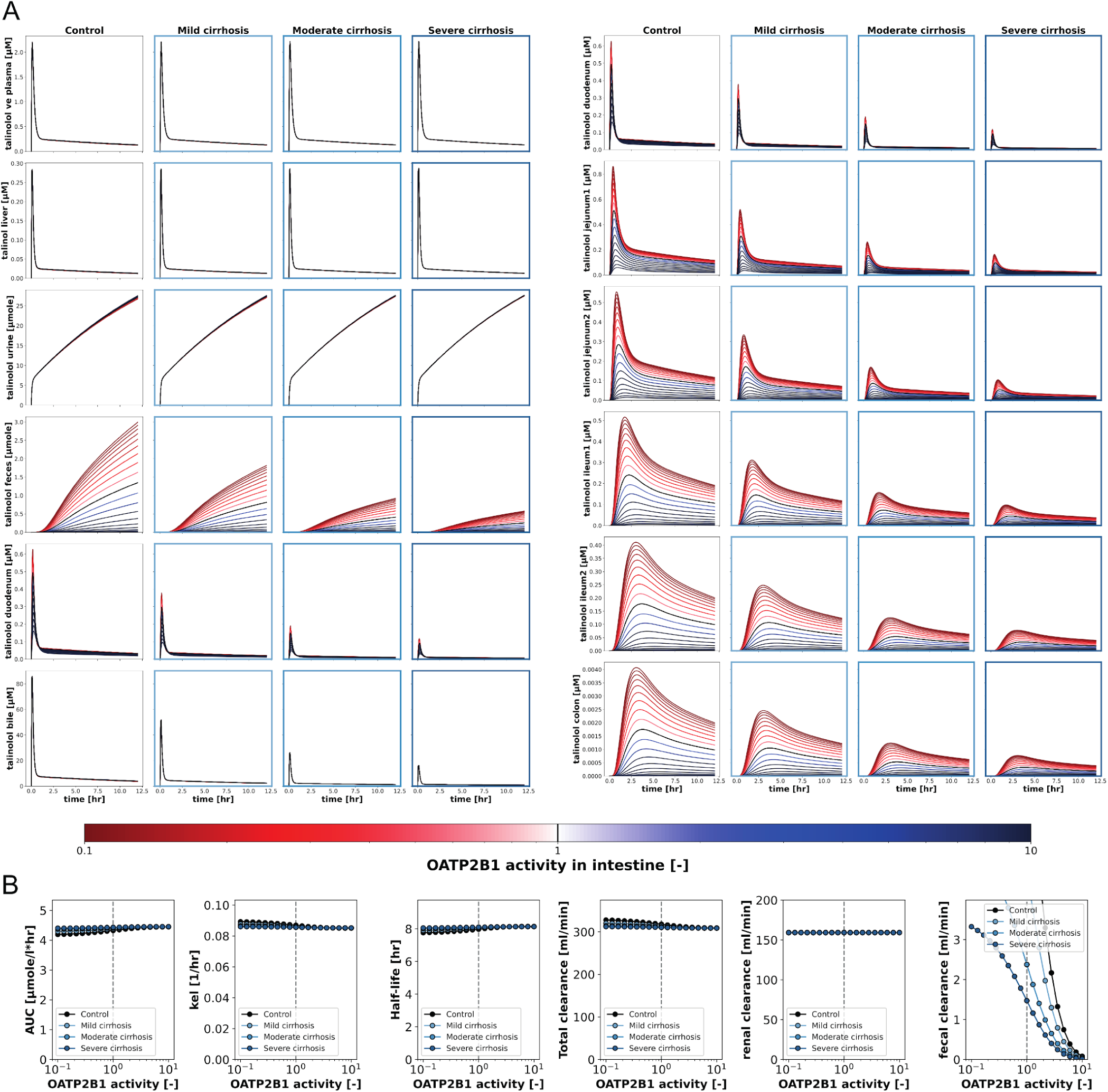
Scan of OATP2B1 activity after intravenous administration of talinolol. 30 mg of talinolol was administered intravenously and OATP1B1 activity was varied as `np.logspace(start=-1, end=10, num=19)` corresponding to a scan of `GU f OATP2B1` from 0.1 to 10. **(A)** Overview of the effects of protein activity on the time course of talinolol in different compartments at different degrees of cirrhosis. Increased and decreased protein activities are indicated by the blue and red gradients, respectively. **(B)** Overview of the effects of protein activity on talinolol pharmacokinetic parameters in different degrees of cirrhosis: Area under the curve (AUC), elimination rate (k_el_), half-life (t_half_) and clearance (renal, faecal and total).

**Figure S8.**
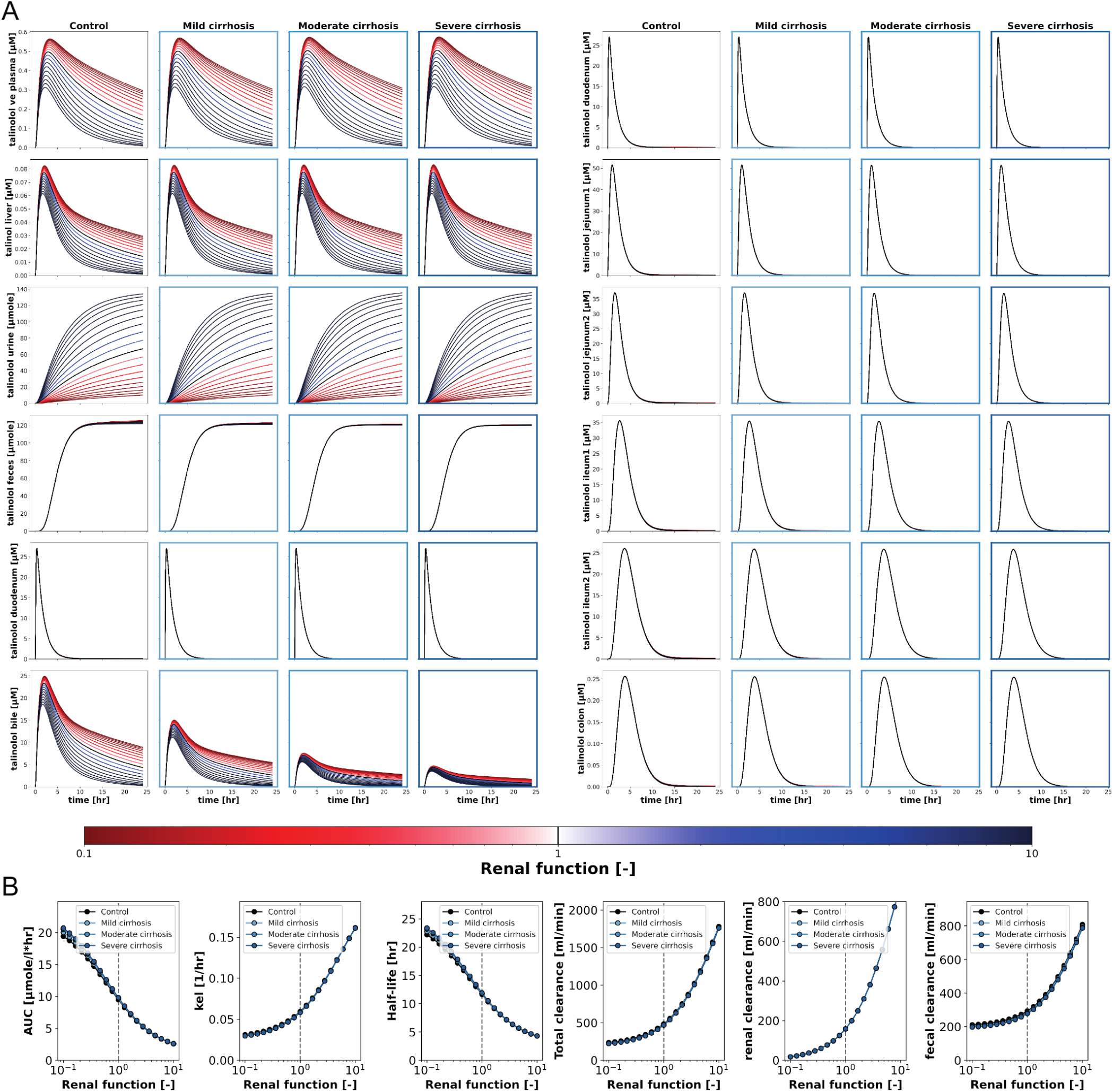
Scan of renal function after oral administration of talinolol. 100 mg of talinolol was administered orally and renal function was varied as `np.logspace(start=-1, end=10, num=19)` corresponding to a scan of `KI f renal function` from 0.1 to 10. **(A)** Overview of the effects of renal function on the time course of talinolol in different compartments at different degrees of cirrhosis. Increased and decreased protein activities are indicated by the blue and red gradients, respectively. **(B)** Overview of the effects of renal function on talinolol pharmacokinetic parameters in different degrees of cirrhosis: Area under the curve (AUC), elimination rate (k_el_), half-life (t_half_) and clearance (renal, faecal and total).

**Figure S9.**
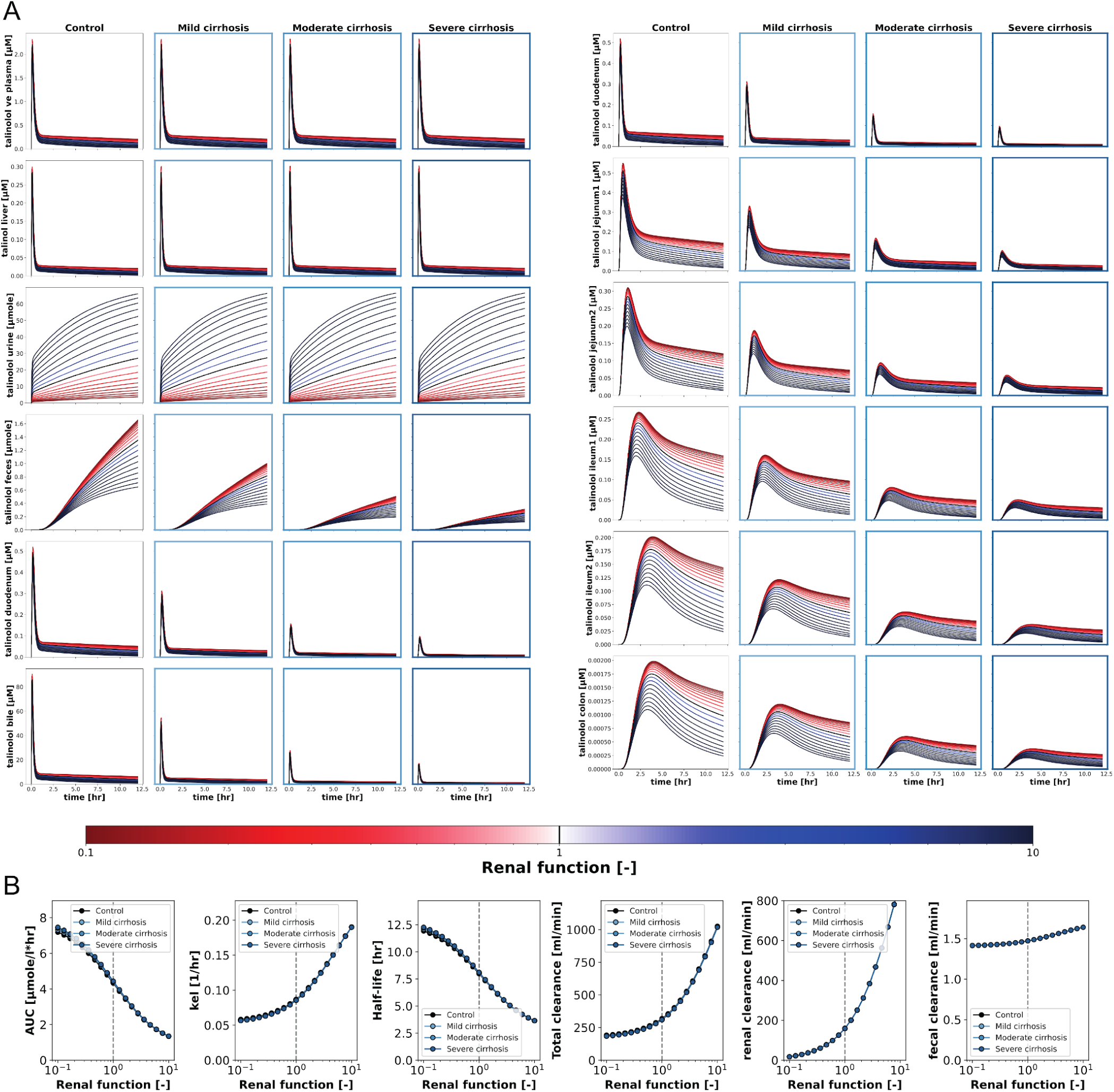
Scan of renal function after intravenous administration of talinolol. 30 mg of talinolol was administered intravenously and renal function was varied as `np.logspace(start=-1, end=10, num=19)` corresponding to a scan of `KI f renal function` from 0.1 to 10. **(A)** Overview of the effects of renal function on the time course of talinolol in different compartments at different degrees of cirrhosis. Increased and decreased protein activities are indicated by the blue and red gradients, respectively. **(B)** Overview of the effects of renal function on talinolol pharmacokinetic parameters in different degrees of cirrhosis: Area under the curve (AUC), elimination rate (k_el_), half-life (t_half_) and clearance (renal, faecal and total).

